# Common nitrification inhibitors exhibit varied physiological mechanisms on an ammonia-oxidizing microorganism

**DOI:** 10.64898/2026.05.10.724060

**Authors:** Dimitrios Dalkidis, Andrea Malits, Melina Kerou, Hadis Sajedi, Leila Afjehi-Sadat, Christa Schleper, Dimitrios G Karpouzas, Evangelia S Papadopoulou, Logan H. Hodgskiss

**Affiliations:** University of Thessaly, Department of Environmental Sciences, Laboratory of Environmental Microbiology and Virology, Larissa, Greece; University of Vienna, Department of Functional and Evolutionary Ecology, Djerassiplatz 1, A-1030 Vienna, Austria; University of Vienna, Research Support Facilities, University Biology Building, Mass spectrometry Unit, Djerassiplatz 1, A-1030 Vienna, Austria; University of Thessaly, Department of Biochemistry and Biotechnology, Laboratory of Plant and Environmental Biotechnology, Larissa, Greece

## Abstract

Microbial ammonia oxidation, the first and rate-limiting step of nitrification, plays a central role in soil nitrogen cycling. It is most relevant in agricultural soils as nitrifiers compete with crops for ammonia-based fertilizers. Therefore, synthetic nitrification inhibitors are widely used alongside fertilizers to reduce the activities of dominant drivers of this process, i.e. ammonia-oxidizing archaea (AOA) and bacteria (AOB). However, the physiological responses of ammonia oxidizers remain poorly resolved. Here the response of the AOA *Nitrososphaera viennensis* to the nitrification inhibitors 3,4-dimethylpyrazole phosphate (DMPP) and allylthiourea (ATU) were investigated using a combination of functional genomics, physiological assays, and relief experiments. The results overturn earlier assumptions that DMPP and ATU act by chelating free copper. Both compounds affected ammonia oxidation and triggered broader shifts in energy metabolism and stress-response pathways, which diverged markedly between the two inhibitors. We propose a competitive inhibition of the ammonia monooxygenase complex with DMPP as it can be alleviated by additional ammonia and elicits activation of urea acquisition, while ATU acted as a non-competitive inhibitor generally inducing quiescence. Both modes of inhibition were associated with clear transcriptomic and proteomic signals that will be advantageous for the identification of mechanisms of other nitrification inhibitors in the future.

Key word: Ammonia-oxidizing archaea, nitrification, nitrification inhibitors, archaea, nitrogen cycle

## Introduction

Since the middle of the 20th century, human activity has significantly intensified its impact on the global nitrogen cycle^1,2^. Due to activities such as burning fossil fuels and producing fertilizers, the amount of fixed nitrogen in Earth’s ecosystems has dramatically increased^3^. Fixed nitrogen is primarily used for fertilization, but as a side effect leads to high rates of nitrification in soils. This reduces crop nitrogen use efficiency (NUE) dramatically^4,5^ and causes eutrophication and nitrogen leaching into the groundwater and coastal zones^6–8^, while also increasing the amount of the potent and long-lived greenhouse gas nitrous oxide (N_2_O) in the atmosphere worldwide^9^. A potential solution to this problem is the inhibition of the activity of nitrification in agricultural systems.

The process of nitrification is carried out by distinct groups of microorganisms. The first step of nitrification is the oxidation of ammonia to hydroxylamine performed by ammonia-oxidizing archaea (AOA), ammonia-oxidizing bacteria (AOB), or complete ammonia-oxidizing (comammox) bacteria, that all utilize the copper-dependent enzyme ammonia monooxygenase (AMO)^10^. The further oxidation to nitrite (NO ^-^) by each of these clades remains incompletely understood^11^. While the enzyme controlling the step following AMO has been identified as hydroxylamine dehydrogenase in AOB^12,13^ and comammox^14,15^, the exact reactions and enzymes involved downstream to AMO are still unknown in AOA^16,17^. Among ammonia-oxidizing microorganisms, AOA often outnumber their bacterial counterparts in soil and play a key role in the global biogeochemical cycling of nitrogen^18–20^. A major advance towards determining their importance and role was the successful isolation of the first AOA from terrestrial environments, *Nitrososphaera viennensis*^21,22^, now taxonomically assigned to the archaeal phylum Thermoproteota (GTDB classification system)^23^. It serves as a model AOA strain for soil systems as it is well studied and characterized at a physiological and genomic level^16,21,22,24–30^. The availability of well-studied model strains that are abundant in soil presents the opportunity to study the inhibition of nitrification at a high level of physiological detail.

In agricultural practice, nitrification inhibition is attempted through the use of nitrification inhibitors (NIs), a group of synthetic chemical compounds that are supposed to target ammonia-oxidizing microorganisms and modulate microbial processes driving ammonia oxidation. Since the early 2000s, the synthetic NI 3,4-dimethylpyrazole phosphate (DMPP) has been commercially used in stabilized nitrogen fertilizers^31^, albeit little is known about its inhibitory mechanism. A recent study showed that DMPP can chelate copper, a metal critical to the functioning of AMO^32^. However, a subsequent study in AOB suggested that its inhibitory effect is not primarily related to its copper chelating activity, but rather to its capacity to interfere directly with AMO in the first step of ammonia oxidation^33^.

Apart from DMPP which is a commercial NI used in agricultural settings, allylthiourea (ATU) is a well-known NI commonly used for research purposes. It is often applied at low concentrations to specifically inhibit AOB but has also been shown to be effective on AOA at higher concentrations^30^. Previous work with ATU has also shown that it specifically targets ammonia oxidization as opposed to hydroxylamine oxidation in AOB^33,34^ and proposed that it acts either as a copper chelator^34–36^ or by inhibiting the transport of ammonium into the cell^37^. However, a general copper chelation mechanism seems unlikely, as no recovery is obtained by AOB when additional copper is added^33,37^. Similar to DMPP, previous work demonstrated that ATU is likely to directly target the AMO as shown in the marine AOA *Nitrosopumilus maritimus*^38^.

Although DMPP and ATU are commonly used for agricultural and research purposes respectively, a deeper understanding of their physiological effects on any nitrifier is still lacking, exemplifying the substantial knowledge gap about nitrification inhibitors and other chemicals widely used in agriculture. This study aims to address this knowledge gap in the model AOA strain *N. viennensis* using an integrative approach combining multi-omics, target specificity investigations, and relief experiments with alternative/excess substrates (ammonium, urea) or key cofactors (copper). DMPP and ATU were chosen due to their similar proposed mechanisms (i.e., copper chelation and/or inhibition of the AMO), but distinct chemical structures that would likely initiate distinct cellular responses detectable by the use of both transcriptomic and proteomic approaches. The study provides previously unknown insights into the mechanisms of ATU and DMPP, and establishes a framework for the testing of novel NIs in the future.

## Results

### Distinct Target Specificity of DMPP and ATU revealed by microrespirometry

The potential target specificity of DMPP and ATU (Fig. 1A) was deduced through tracking the oxygen consumption of *N. viennensis* cells in a microrespiratory chamber. High concentrations of both DMPP (5 mM) and ATU (3 mM) showed substantial reduction of oxygen consumption as an indication for the inhibition of physiological activity, when compared with control cultures in the presence of ammonium (Fig. 1B). A higher DMPP concentration of 7 mM was also tested in an attempt to achieve full inhibition with similar results (Fig. S1). Full inhibition by DMPP by using even higher concentrations was not possible due to acidification of the medium, even in the presence of additional HEPES buffer (50 mM). When amended with hydroxylamine, ATU (3 mM) showed no inhibitory effect while DMPP (5 mM) showed a slight reduction in the oxygen consumption rate compared to hydroxylamine controls (Fig. 1C).

**Figure 1.**
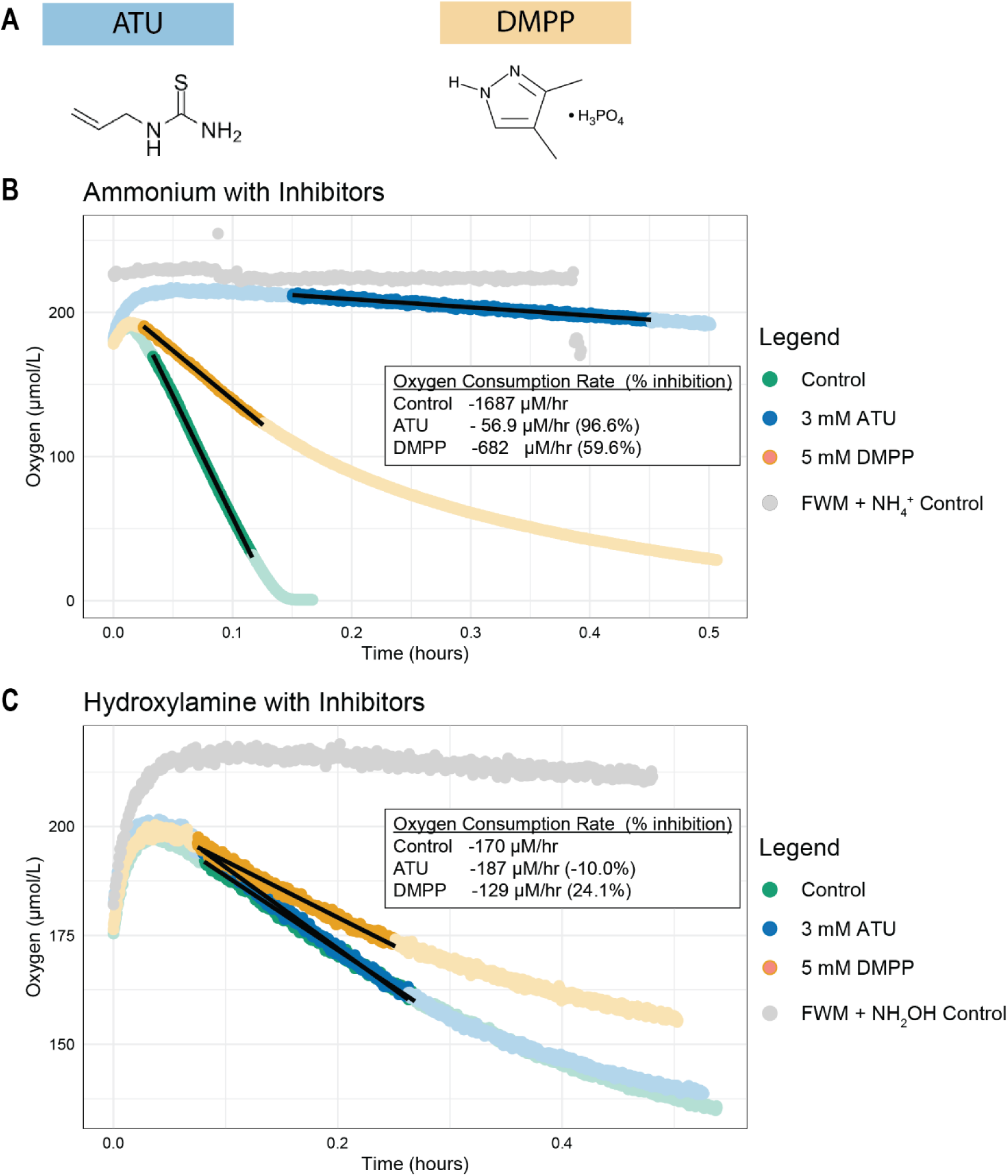
Micro-respiration assay on *N. viennensis*. **A)** Chemical structures of DMPP and ATU. **B)** *N. viennensis* exposed to ATU (3 mM) or DMPP (5 mM) in the presence of ammonia (200 μM) as substrate. **C)** *N. viennensis* exposed to ATU (3 mM) or DMPP (5 mM) in the presence of hydroxylamine (200 μM) as substrate. Darker regions indicate the portion of the curve used to calculate the rate of oxygen consumption. Black lines represent the calculated slope.

The inhibition pattern of ATU indicates that it is effective only in the presence of ammonium. While DMPP had a much stronger effect in the presence of ammonium (59.6% inhibition), there was still a mild inhibitory effect observed in the presence of hydroxylamine (24.1% inhibition) suggesting that DMPP is also interfering with the archaeal ammonia oxidation process downstream of the initial conversion of ammonia to hydroxylamine.

### DMPP addition shows immediate transcriptional responses

The determination of the appropriate time for harvesting to capture effects in both transcriptome and proteome was crucial to further investigate the effect of NIs on *N. viennensis*. Cells harvested at different time points following DMPP addition were therefore evaluated compared to cells from a control culture without DMPP to determine the dynamics of transcription and translation within *N. viennensis* (Fig. 2A). Sequencing of rRNA-depleted samples resulted in an average of 26.45 million paired end reads per sample while protein sequences using mass spectrometry resulted in a proteome coverage of 32.38% (1,032 of 3,187 predicted proteins). Principal component analysis (PCA) of normalized RNA sequences (rlog of transcripts per million (TPM)) revealed that transcriptomic responses were detectable as early as 0.5 hours following DMPP addition, as indicated by clear separation from control samples (Fig. 2B). In contrast, a PCA of the label-free quantification (LFQ) protein values after filtering, imputation, and log_2_ transformation revealed that proteomic responses were only evident after at least 6 hours, with substantial overlap between treated and control samples observed at the earlier time points of 0.5, 1 and 3 hours (Fig. 2C).

**Figure 2.**
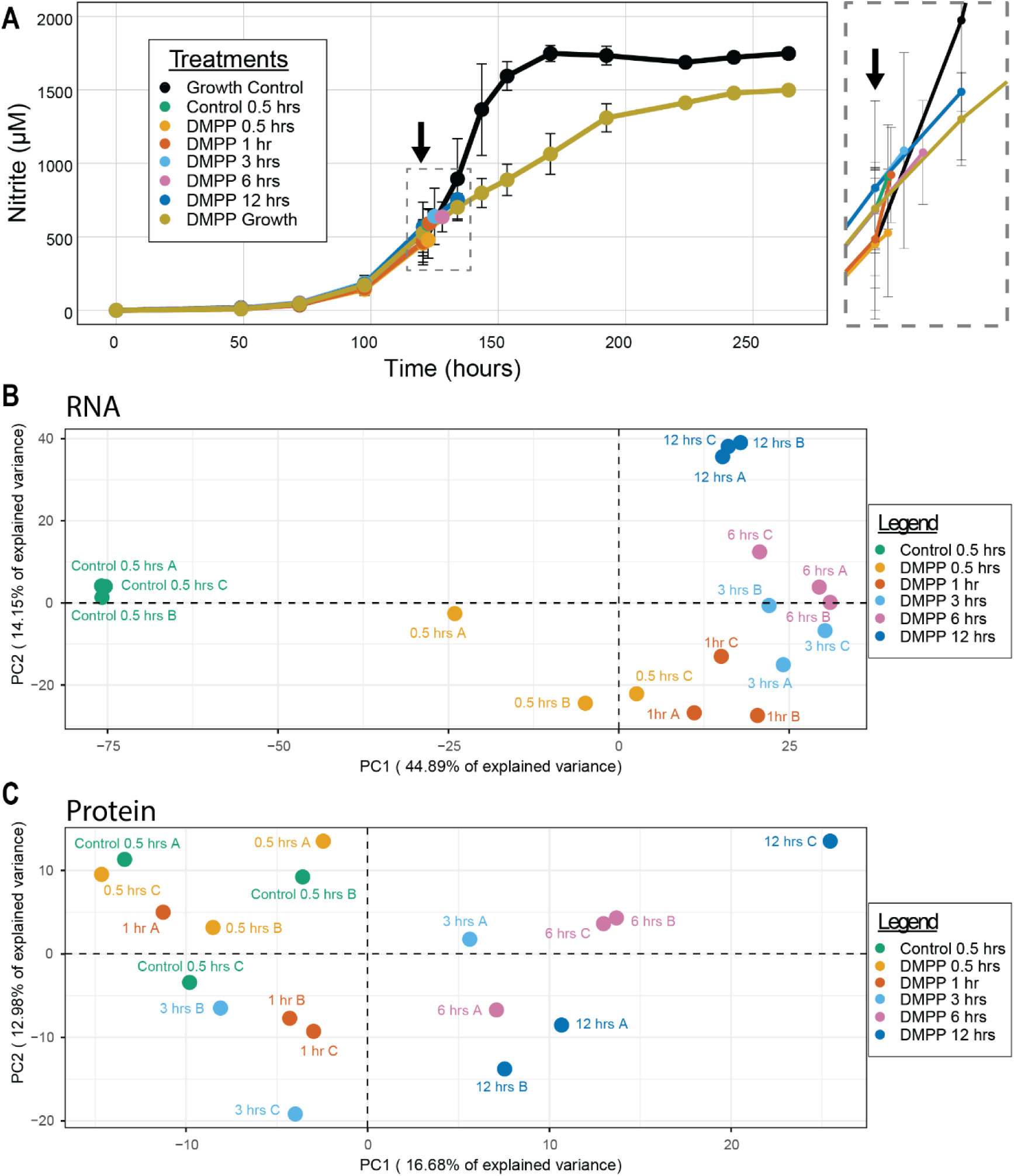
Time-series experiment on *N. viennensis* with 2 mM DMPP. **A)** The effect of 2 mM DMPP on *N. viennensis* activity. Error bars represent standard deviation of the mean (triplicates). Black arrows indicate the time of DMPP addition. Zoomed panel shows the nitrite level for each condition when DMPP was added and when the cultures were harvested. **B)** Principal component analysis including all detected genes (transcripts per million (TPM)) from transcriptome analysis. The percentage of explained variance in the first two principal components (PCs) is shown. **C)** Principal component analysis of all detected protein values (normalized and imputed label-free quantification (LFQ) values) from proteomic analysis.

Genes exhibiting significantly different expression patterns in the presence or absence of DMPP (adjusted *P* value < 0.001) were grouped into six clusters based on similar expression dynamics and visualized in a heatmap (Fig. 3A). These clusters revealed distinct transcriptional response patterns, including genes with decreasing expression (Cluster 1, 587 genes; Cluster 5, 136 genes), increasing expression (Cluster 3, 258 genes; Cluster 6, 522 genes), or transient expression patterns (down then up, Cluster 2, 202 genes; up then down, Cluster 4, 181 genes). Functional enrichment analysis was conducted for each cluster to assess whether specific archaeal clusters of orthologous gene (arCOG) categories^39^ were associated with shared expression patterns (Fig. 3B). Cluster 1, representing immediate down-regulation, was enriched with genes from the following arCOG categories: translation, ribosomal structure and biogenesis; nucleotide transport and metabolism; lipid transport and metabolism; energy production and conversion; coenzyme transport and metabolism; and amino acid transport and metabolism. In contrast, cluster 3, represented by genes whose abundance steadily increased over time, was enriched for functions related to transcription initiation and regulation, signal transduction, post-translational modification, protein turnover, chaperones and cell motility.

**Figure 3.**
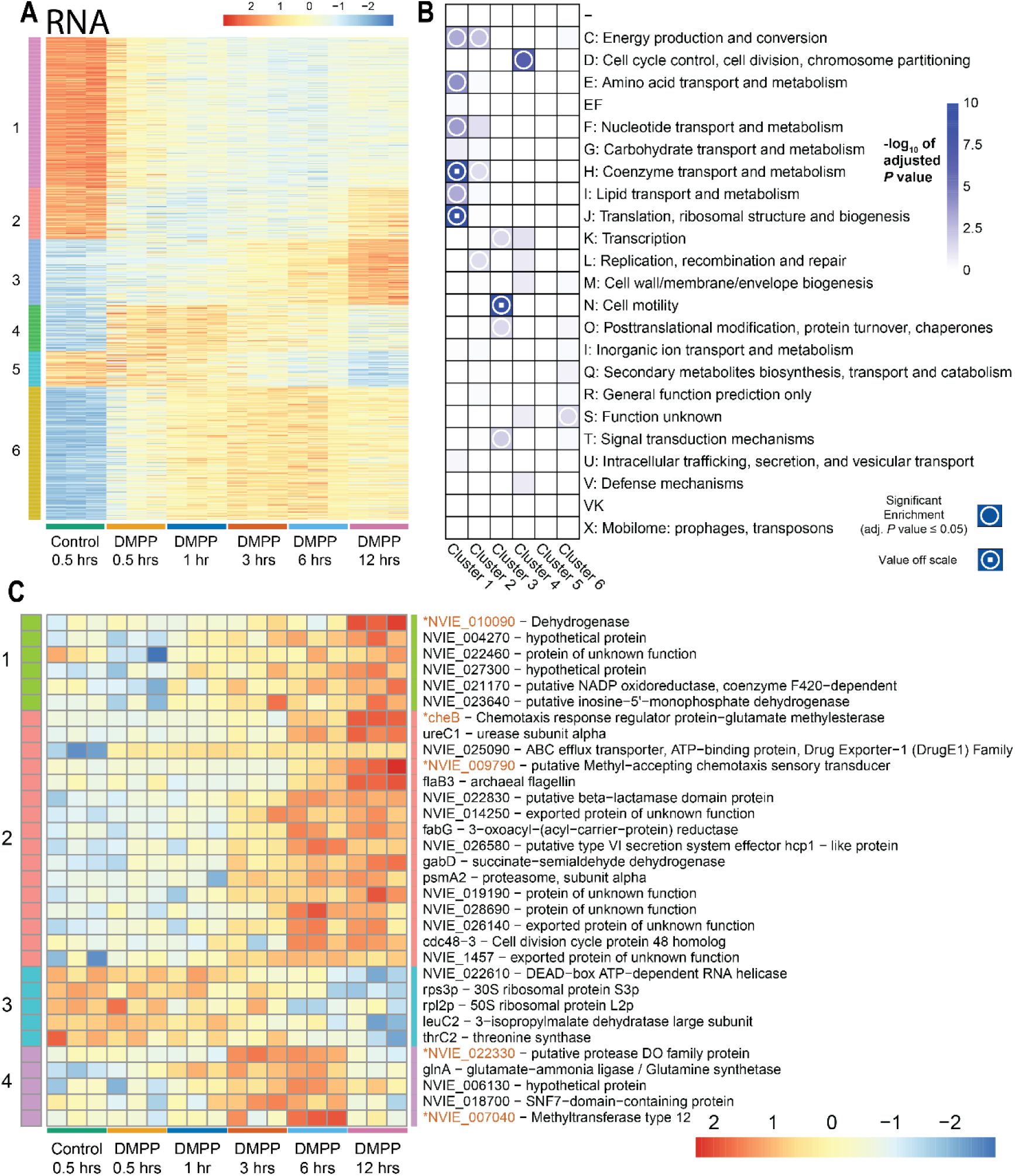
Clustered heatmap of significantly changing transcripts and proteins over a 12-hour time period following exposure to 2 mM DMPP. **A)** Transcriptomic heatmap of genes identified as significantly differentially expressed (adjusted *P* < 0.001) using a DESeq2 Likelihood Ratio Test (LRT). Expression values (transcripts per million) were rlog-transformed, centered, and scaled before plotting. Columns are ordered chronologically (Control to 12h post-DMPP treatment). **B)** Enrichment analysis of differentially expressed genes based on arCOGs in every cluster from transcriptomic heatmap. Boxes with a white circle indicate arCOG categories that are enriched in their respective cluster based on a hypergeometric test. Boxes with a white circle and a white square indicate values off scale. EF represents a combined category of amino transport and metabolism (E) and nucleotide transport and metabolism (F). VK represents a combined category of defense mechanisms (V) and transcription (K). **C)** Proteomic heatmap of proteins identified as differentially regulated (adjusted *P* value < 0.05) using analysis of variance (ANOVA). Data represent outlier-corrected, normalized, and imputed log_2_-transformed label free-quantification (LFQ) intensities. Data were centered and scaled before plotting. Columns are ordered chronologically (Control to 12h post-DMPP treatment). Proteins in orange marked with an * are proteins where 67% or more of replicates relied on imputation due to missing values.

Unlike transcription, translational responses showed a delayed response after the addition of DMPP. Proteins showing significant changes over time were identified using analysis of variance (ANOVA, adjusted *P* value < 0.05). Thirty-two proteins were identified as significantly changing with time under DMPP exposure and were grouped into four clusters and visualized in a heatmap (Fig. 3C). Consistent with transcriptomic trends, protein abundance either increased (Cluster 1, 6 proteins; Cluster 2, 16 proteins), decreased (Cluster 3, 5 proteins), or showed a transient expression pattern (up then down, Cluster 4, 5 proteins) as exposure time progressed. Enrichment analysis was not performed due to the low number of statistically significant proteins in each cluster.

From the time-series analysis, it was concluded that an optimal timepoint for analyzing the transcriptome and proteome simultaneously was at 9 hours after addition of the inhibitor. This represents ∼70% of the generation time of *N. viennensis* and would reliably capture changes in the proteome while transcriptomic responses are still maintained, albeit a stronger transcriptional response was observed in earlier timepoints.

### Comparative omics reveal differences in responses to DMPP versus ATU

Cultures of *N. viennensis* were treated with 2 mM DMPP or 0.5 mM ATU and harvested after 9 hours to compare the physiological effects of the two nitrification inhibitors (Fig. 4A). Comparative transcriptomic analysis of cells harvested 9 hours following addition of ATU or DMPP yielded an average of 12.84 million paired-end reads per sample, while proteomics led to a proteome coverage of 45.06% (1,436 of 3,187 predicted proteins) using an improved protocol (see materials and methods). This data showed a high degree of overlap between ATU and DMPP, but also distinct physiological differences (Fig. 4B-E).

**Figure 4.**
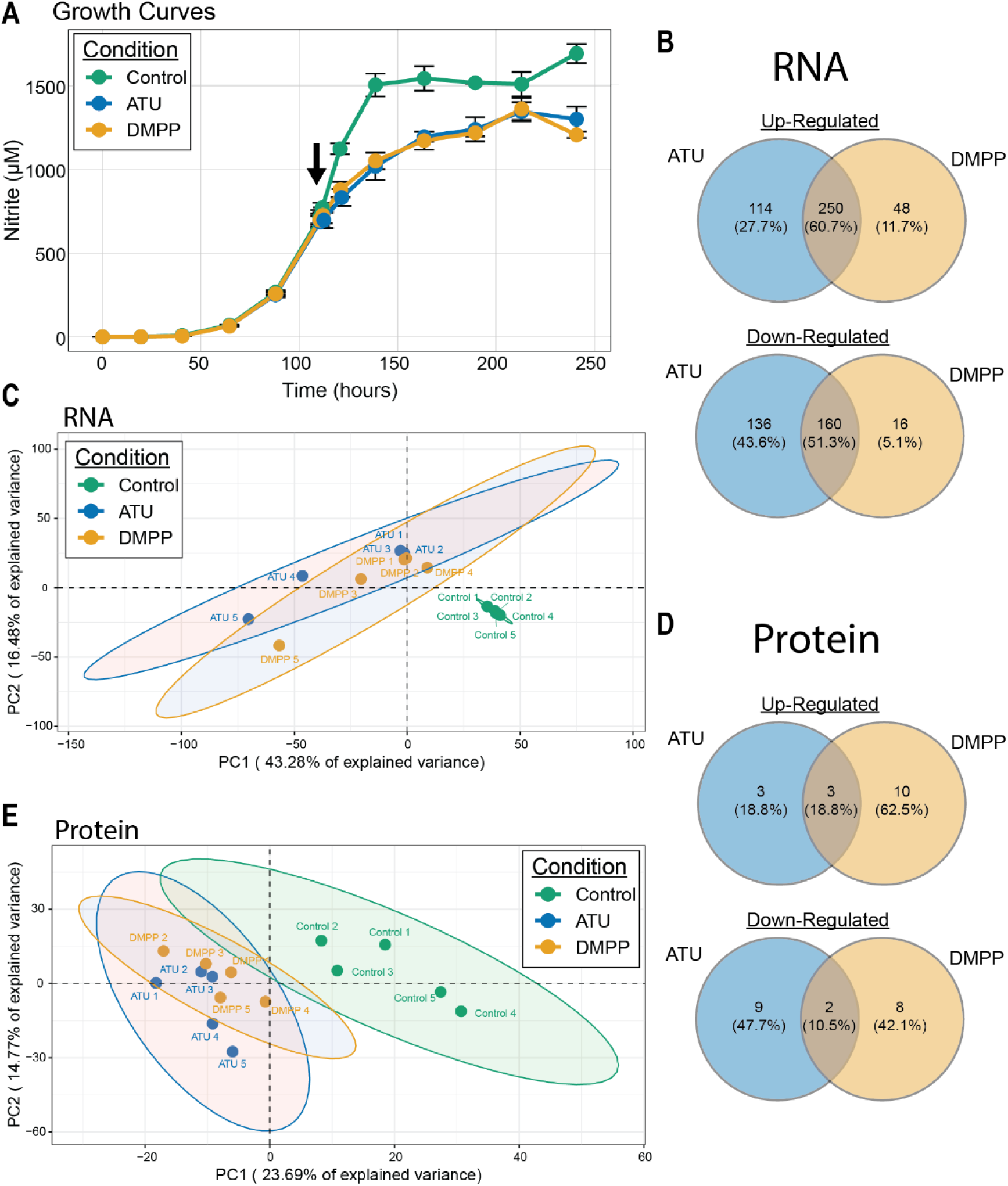
Transcriptomic and proteomic response of *N. viennensis* to 2 mM DMPP and 0.5 mM ATU. **A)** The effect of 2 mM DMPP and 0.5 mM ATU on *N. viennensis* growth. Error bars represent standard deviation of the mean. Black arrow represents the time of DMPP or ATU addition. **B)** Venn diagrams of the common up-regulated and down-regulated genes between DMPP and ATU treatments. **C)** Principal component analysis including all detected genes (transcripts per million (TPM)) from transcriptome analysis. The percentage of explained variance in the first two principal components (PCs) is shown. **D)** Venn diagrams of the common up-regulated and down-regulated proteins between DMPP and ATU treatments. **E)** Principal component analysis of all detected protein values (normalized and imputed label-free quantification (LFQ) values) from proteomic analysis.

Overall, ATU elicited a more severe physiological response, characterized by extensive transcriptional repression and down-regulation of pathways associated with energy metabolism. Transcripts that were specifically down-regulated in ATU but not DMPP included all subunits of the terminal oxidase, two multi-copper oxidases (MCO4a and MCO4b), nitrite reductase (*nirK*), and two blue copper proteins suspected to be involved in electron transfer (NVIE_029580 and NVIE_005340) (Fig. 5). A strong down-regulation of the terminal oxidase was surprising as this is the only suspected contributor to the formation of a proton motive force in AOA. MCO4a^40^ and *nirK*^41^ are also suspected of playing stress-relieving roles in electron transfer, and their down-regulation suggests critical issues in cellular metabolism. Additional evidence for severe impairment can also be seen in the transcripts of the cell division machinery of AOA (*cdvA*, *cdvB*)^42^ that are primarily down-regulated under ATU stress. At a proteomic level, the most down-regulated protein is the large subunit of the DNA polymerase family D (annotated as PolC). While there are few up-regulated genes or proteins that help to define the response of ATU, an exception is the translation initiation factor protein eIF-2B. In this case, the protein was fully detected in all ATU replicates but had to be imputed in all controls and DMPP due to low abundance. Oddly, this is the only translation initiation factor affected.

**Figure 5.**
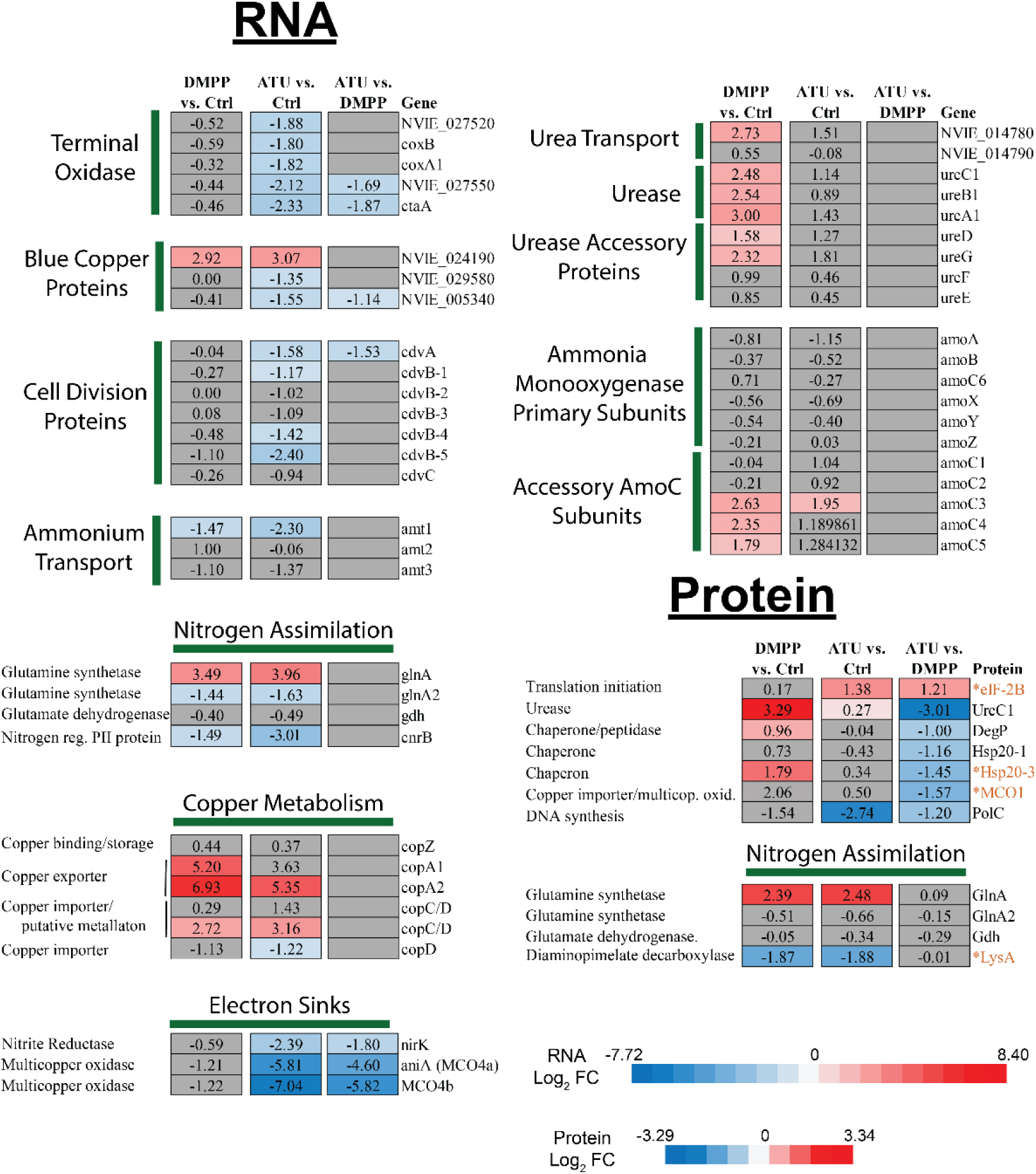
Selected differentially expressed genes and regulated proteins (univariate analysis). Genes were considered to be differentially expressed with an adjusted *P* value < 0.001 and a log_2_ fold change >1 or < -1. Proteins were considered differentially regulated with an adjusted *P* value < 0.05.). Proteins in orange marked with an * are proteins where all replicates in one condition relied on imputation due to missing values. Full gene and protein annotations can be found in Dataset S1.

In stark contrast to ATU, stress initiated by DMPP was largely defined by the up-regulation of genes and proteins responding to an energy limitation signal. At the transcript level, this was captured by the up-regulation of all primary urease subunits (*ureABC1*), two urea accessory proteins (*ureDG*), and the adjacent urea transporter (NVIE_014780) (Fig. 5). It was also reflected in the up-regulation of two auxiliary *amoC* subunits (*amoC4* and *amoC5*), as well as an increase in gene expression of phosphate transporters, a critical molecule for both energetics and DNA. In strong support of the transcriptomic results, the highest up-regulated protein was UreC1, the subunit containing the active site of the urease complex. Unique to DMPP was also the up-regulation of a heat shock protein (Hsp20-3) previously implicated in energy stress in AOA^43^, and a protease that has also been shown to have chaperone-like functions (DegP)^44^.

Two additional comparisons were used to further explore differences between the ATU and DMPP treated cultures that were not initially obvious: 1) a direct univariate approach comparing the response of ATU to DMPP to identify statistically significant differences in genes and proteins (as done between each treatment and control) (Fig. 5), and 2) a sparse partial least squares discriminant analysis (sPLS-DA) multivariate approach to detect underlying genes and proteins that can distinguish between conditions but may not be highlighted when testing each gene or protein individually (Fig. 6). In sPLS-DA analysis, a subset of variables (genes or proteins) is selected that maximizes the separation of each condition (see supplementary material and methods).

**Figure 6.**
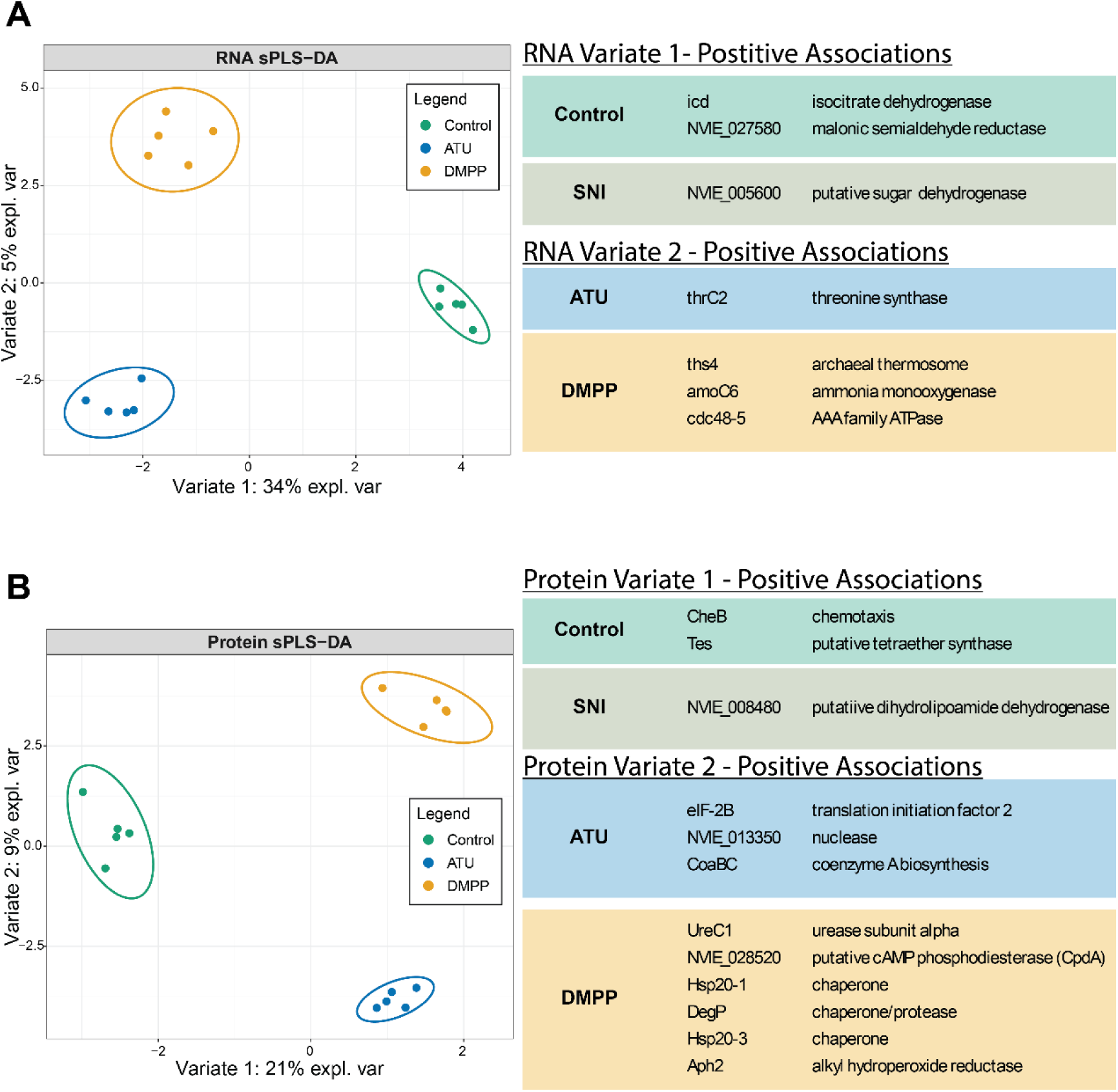
Multivariate sparse partial least squares discriminant analysis (sPLS-DA). **A)** sPLS-DA analysis using 10 genes that best separate along variate 1 and 20 genes that best separate along variate 2 (transcripts per million (TPM)) from transcriptome analysis. The percentage of explained for each variate is also shown. Selected genes positively associated with different conditions are shown to the right. **B)** sPLS-DA analysis using 5 proteins that best separate along variate 1 and 25 proteins that best separate along variate 2 (normalized and imputed label-free quantification (LFQ)) from proteome analysis. The percentage of explained variance for each variate is also shown. Selected proteins positively associated with different conditions are shown to the right. Methods for selecting number of genes and proteins can be found in Supplementary Materials and Methods. Full lists of genes and proteins selected for plotting can be found in Dataset S1.

At the RNA level, the univariate approach (Fig. 5) emphasized the distinct differences outlined above. At the protein level, a membrane bound multicopper oxidase previously speculated as important for copper import (MCO1)^25,45^ was identified as being more highly abundant in DMPP than in ATU. While present in all five replicates of DMPP, MCO1 was not found in any ATU replicates and only two control replicates and therefore interpretation was reliant on a large amount of imputation, possibly explaining the lack of a strong response (Fig. S2). In both the RNA and protein datasets, the multivariate approach using sPLS-DA analysis was able to separate the control from NI treatments along variate-1 and was able to separate ATU from DMPP along variate-2 using a selected subset of genes or proteins. The subset of genes or proteins for each plot is positively associated with a particular condition, i.e., they could be categorized by which condition they are likely to be more abundant. The analysis identified *amoC6*, the primary amoC subunit, and an archaeal thermosome as being positively associated with DMPP (more abundant in DMPP than in ATU) at the RNA level, and an additional heat shock protein, as well as the previously identified UreC1, as positively associated with DMPP (more abundant in DMPP than in ATU) at the protein level (Fig. 6). The increased abundance of *amoC6* and thermosome transcripts in DMPP compared to ATU, and the increased abundance of an additional heat shock protein, supports the observation that DMPP stress is causing an energy limitation signal in a way that is not observed during ATU stress.

Despite these clear differences, a large overlap in responses was also observed. Many highly up-regulated transcripts of unknown function were found in both DMPP and ATU (Fig. S3, Dataset S2). Up-regulated genes of particular interest included a copper exporter (copA2), one copy of copC/D putatively involved in copper metabolism (NVIE_014310), the auxiliary *amoC3* subunit, one copy of glutamine synthase (*glnA*), and a blue copper protein assumed to transfer electrons during ammonia oxidation (NVIE_024190) (Fig. 5). Both treatments caused a transcriptional down-regulation of Complex I and ATP synthase of the electron transport chain (Fig. S4) as well as the second copy of glutamine synthase (*glnA2*). Surprisingly, both treatments also caused a down-regulation of one ammonium transporter (*amt1*) and a PII nitrogen regulatory protein (NVIE_013340).

At the protein level, many of the commonly up- or down-regulated proteins were of unknown function. A strong exception is the up-regulation of the glutamine synthetase, GlnA. The expression of the alternative glutamine synthetase, GlnA2 remained constant in both cases (Fig. S5). Another exception is the down-regulation of LysA putatively involved in ammonium metabolism within the cell (Fig. 5). However, the extent of down-regulation is only estimated as LysA was only detected in control cultures and all values in DMPP and ATU conditions were estimated by imputation (Fig. S6). While both DMPP and ATU resulted in down-regulation of the chemotaxis protein (CheB) (Fig. S6, Dataset S3), these results are counter to what was observed in the DMPP time-series experiment and should be treated with caution (see supplementary discussion). However, CheB abundances had to be estimated by imputation for all replicates of DMPP and ATU as well as for all control and early time points of the time-series (Fig. S6), thereby lowering the confidence of this response.

### Relief Experiments

The exclusive or strong inhibitory effect of ATU and DMPP, respectively, in the presence of ammonium but not hydroxylamine along with results from the functional omics provided a basis for testing possible relief mechanisms for each inhibitor. In this framework, liquid batch cultures were amended with copper (3000 µM), trace elements (10x), ammonium (5 mM), or urea (1 mM and 2.5 mM) to explore the potential of NIs for copper chelation, alternative metal chelation, competitive inhibition of ammonia, and the blocking of ammonia transporters respectively.

The addition of copper or trace elements to fresh water medium (FWM) medium did not relieve inhibition in ATU or DMPP rendering the previously proposed mechanism of copper or metal chelation unlikely (Fig. 7AB). The concentration of additional copper and other metals was far below the concentration of DMPP or ATU to avoid toxicity effects from the metals to the AOA strain^46^. While it is possible that more copper or trace elements could be added, this is unlikely to be effective as lower concentrations of DMPP and ATU do not inhibit *N. viennensis*, but are still orders of magnitude higher than copper or trace metal concentrations. For these reasons, metal chelation was discounted as a mechanism of action.

**Figure 7.**
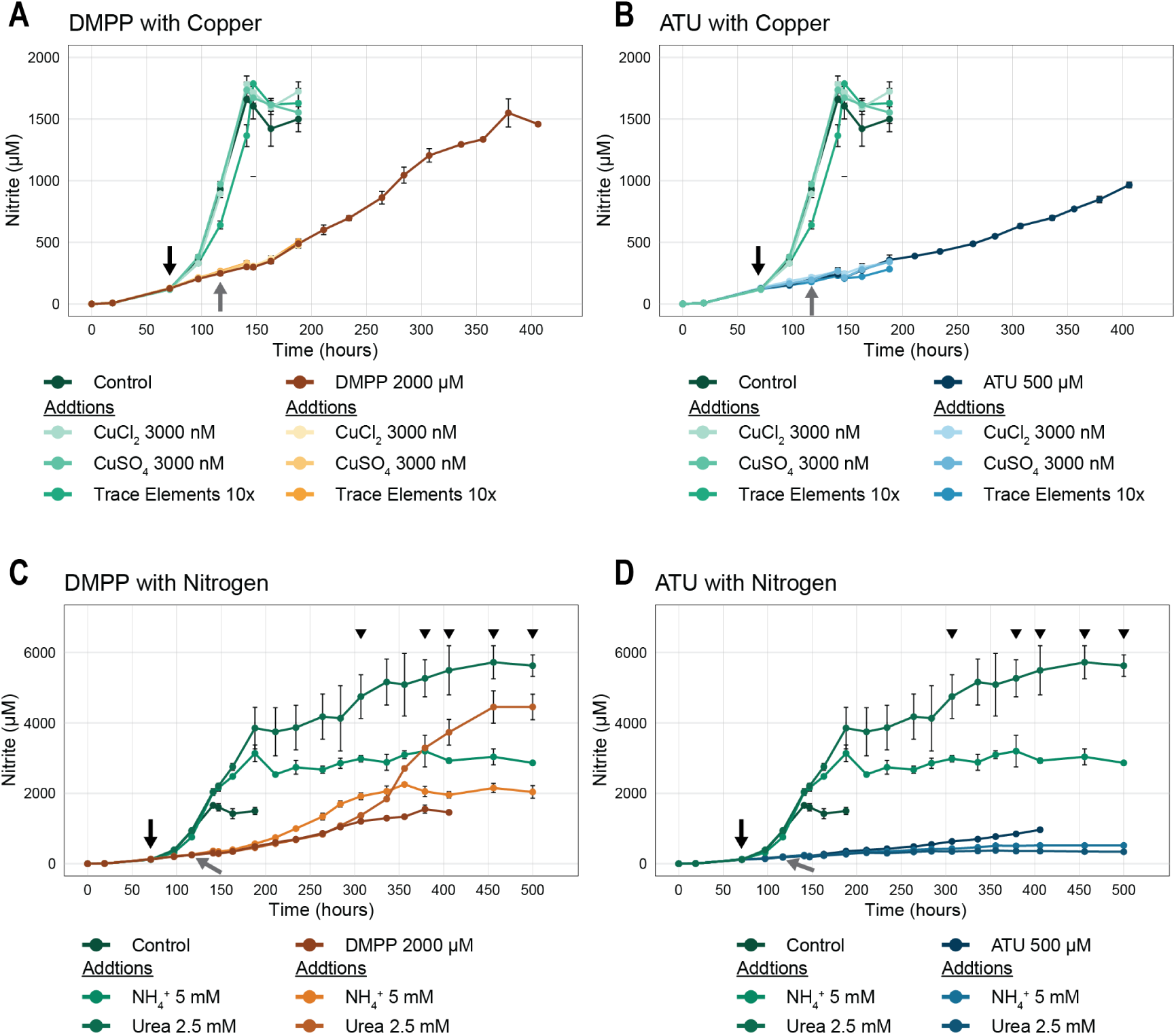
Stress relief experiments on *N. viennensis*. A and. **B)** Additions of copper and trace elements to cultures treated with 2 mM DMPP **(A)** or 0.5 mM ATU **(B)**. **C and D)** Additions of ammonium and urea to cultures treated with 2 mM DMPP **(C)** or 0.5 mM ATU **(D)**. Error bars represent standard deviations of the mean (triplicates). Black arrows indicate the time of addition of DMPP or ATU to treated cultures and addition of copper, trace elements, ammonium, or urea to control cultures. Grey arrows represent the time of additions of copper, trace elements, ammonium, or urea to cultures treated with DMPP or ATU. Black triangles represent time points where ammonium measurements were taken for cultures that received additional ammonium or urea. Ammonium measurements can be found in Fig. S8.

Consistent with the multi-omics results, the addition of larger concentrations of urea (2.5 mM) or ammonium (5 mM), but not lower concentrations of 1 mM urea (Fig. S7), resulted in the relief of stress in DMPP treated cultures but not in ATU cultures (Fig. 7 CD). This is strongly suggestive of a competitive inhibition of DMPP at the active site while ATU is inhibiting in a distinct, non-competitive fashion. Growth after addition of urea in DMPP cultures was delayed by approximately 120 hours compared to ammonium cultures, likely due to the needed activation of urease. In both ATU and DMPP treated cultures, ammonium measurements verified that urea was successfully converted to ammonium (Fig. S8). These results indicate that neither DMPP nor ATU inhibit urease or urea transporters. Moreover, the relief observed with urea would bypass any interference of ammonium transporters by either inhibitor, and therefore this mechanism of action was also discounted.

To verify these results, additional ammonium or copper were also added to microrespiratory chambers along with DMPP and ATU. In support of the liquid batch experiments, additional ammonium was able to relieve stress from DMPP, but not ATU (Fig S1). In the case of both DMPP and ATU, additional copper chloride (5 µM) showed no effect on DMPP or ATU stress when added with ammonium.

Since DMPP exhibited slight inhibition when supplied with hydroxylamine, additional copper chloride (5 µM) was also tested under these conditions. While a slight reduction in the amount of inhibition was observed, the additional copper was not able to fully relieve the slight inhibition caused by 7 mM DMPP when added with 200 µM hydroxylamine (Fig S1).

To test if addition of DMPP or ATU is lethal to cells of *N. viennensis,* washed cell pellets treated with DMPP or ATU were used for inoculation of fresh cultures. In both cases, cultures were able to grow after an initial lag phase compared to control cultures (Fig. S9).

## Discussion

The integrative use of functional omics and physiological tests was able to clearly differentiate the effects of two NIs on the ammonia-oxidizing archaeon *Nitrososphaera viennensis*. Based on the results from microrespiration assays, both DMPP and ATU showed high inhibition levels in the presence of ammonium but not hydroxylamine. While some inhibition was still observed with DMPP in the presence of hydroxylamine, the primary target was clearly before this step. These results indicate that the target of both NIs should either be the AMO complex itself or ammonium transporters as this process has been shown to be transport-dependent in AOA^47^. However, the ability of urea to be transported into the cell and hydrolyzed to ammonia in both DMPP and ATU treated cells eliminates the possibility of transport being the target, leaving the primary candidate target as the AMO complex.

### Inhibitors do not function through the chelation of copper

The functioning of the AMO complex presumably relies on the input of several substrates and co-factors including copper, quinones, oxygen, and ammonia, any of which could be affected by NIs. In the case of DMPP and ATU, the chelation of copper ions has been a proposed mechanism and, for DMPP, was confirmed to be possible using X-ray crystallography^32^. However, this mechanism of action for both ATU and DMPP has been called into question as recovery in AOB has not been observed with copper supplementation of ATU-treated cell^33,37^ and other copper-dependent organisms in soil (i.e., denitrifiers) do not appear to be inhibited by DMPP^48–50^. Copper chelation by ATU was speculated to be the cause of lower intracellular copper concentrations of the methanotroph *Methylosinus trichosporium* OB3b that relies on the homologous copper-containing particulate methane monooxygenase (pMMO)^51^. However, it is also possible that the inactivation of pMMO by ATU negated the need to import copper as *M. trichosporium* can switch to the soluble iron-dependent monooxygenase, an option that is not available in ammonia oxidizers^51^.

In agreement with most previous studies, the functional omics presented here do not support copper chelation as a mode of action for either NI. Both NIs caused an up-regulation of some, but not all presumed copper metabolism genes. The lack of a strong response in the presumed copper importer MCO1^25,45^ was particularly surprising. While this protein showed a stronger response to DMPP at the protein level, and was also seen initially up-regulated in the time-series experiment (Fig. S2), the response was not sustained in a manner indicative of intense limitation of copper. Furthermore, when compared to previous transcriptomic studies of copper limited *N. viennensis*^25^, there is a lack of similarity in the top up- or down-regulated genes (Fig. S10). However, it should be noted that the conditions that led to copper limitation in Reyes et al.^25^ (extended copper limitation with 1 µM of TETA) vs. those implied here (short term stress with 2 mM DMPP or 0.5 mM of ATU) may be difficult to compare directly. Beyond the functional omics, there was also a lack of evidence of copper limitation as no response was observed in either NI when 3 µM of copper (300x normal concentrations) was added to liquid cultures. This concentration is far below the level of DMPP or ATU, even if DMPP chelates in a 4:1 (DMPP:Cu) ratio^32^, which could be argued to be the reason for the lack of a relief response. However, if chelation-inhibition is based on stoichiometric ratios, both inhibitors should inhibit AOA far below the values presented here as the typical copper concentration in FWM medium is 10 nM. It is much more likely that chelation depends on the stability and equilibrium of the formed complex. This was demonstrated in *N. viennensis* previously when copper stress initiated with 1 µM of the copper chelator TETA was relieved by a mere 5 nM of additional copper^29^. For these reasons, it was concluded that chelation of free copper is not the primary mechanism for DMPP or ATU.

### Both nitrification inhibitors elicit an energy limitation response

Beyond copper metabolism, the functional omics coupled with the corresponding growth patterns showed that both NIs induced an energy limitation response. This can be seen not only in the inhibited nitrification, but also in the down-regulation of Complex I and the ATP synthase (Fig. S4). This energy limitation was also reflected in the intracellular nitrogen metabolism through the strong transcriptional up-regulation of *glnA* and down-regulation of *glnA2*, emphasized by the high log_2_ fold change of GlnA at the proteomic level (Fig. S5). The genes *glnA* and *glnA2* code for glutamine synthetase, a key part of ammonia assimilation in AOA^16^. The switch observed here is likely indicative of differing enzymatic and affinity properties of the two copies. In support of a modulation of intracellular nitrogen metabolism, both treatments also exhibited a strong down-regulation of diaminopimelate decarboxylase (LysA), a protein that has previously been observed to be responsive to ammonium levels in *Corynebacterium glutamicum*^52^. Surprisingly, both treatments also down-regulated transcripts of *amt1*, a primary ammonium transporter assumed to exhibit a high affinity for ammonium^53^. While all of these responses indicate that both NIs are strongly affecting the energy status of the cells, the metabolic response beyond this initial characterization becomes strikingly unique.

### Indications for competitive inhibition of DMPP at the active site of ammonia monooxygenase

The up-regulation of urease genes (primary & accessory subunits, and transporter), multiple *amoC* subunits, and chaperone proteins is indicative of a cell responding to stress in its environment by adapting its metabolism. This is fully supported by the recovery of DMPP-treated cells when supplied with high concentrations of ammonia or urea for growth. The relief of DMPP stress by the substrate ammonia is strongly suggestive of competitive inhibition at the active site of the AMO complex.

While this clarifies the primary mechanism of DMPP, it is unable to explain the slight inhibition observed to occur after the AMO complex in the microrespiration experiments. While DMPP could still be chelating copper, the addition of free copper was unable to relieve this inhibition in live cultures (Fig. 7) or in microrespiration experiment with 200 µM ammonium and an addition of 5 µM copper (500x standard conditions). While some relief was observed with 200 µM of hydroxylamine and 5 µM of copper chloride (Fig. S1), the DMPP stress was not fully resolved. One possible explanation could be the interference of DMPP with electron transfer processes that are mediated in AOA by blue copper proteins. While free copper may not be a primary target, there could potentially be interference with protein-bound copper as electrons are transferred from hydroxylamine to the necessary electron carriers. The slight relief by copper in the presence of hydroxylamine and the increased abundance of the presumed copper transporter MCO1 at the protein level in DMPP when compared to ATU are suggestive of a role involving copper downstream of the AMO complex. Nevertheless, the primary mechanism of DMPP in *N. viennensis* is clearly linked to interference with the AMO complex in a manner that is not dependent on copper.

### Non-competitive inhibition by ATU

Conversely to the observations with DMPP, the severe transcriptional down-regulation of the terminal oxidase, cell division genes and genes that help to regulate electron flow (blue copper proteins, *nirK*^41^, MCO4a^40^ and/or depend on electron input (MCO4b), as well as the decrease in DNA polymerase protein levels and the lack of chaperone proteins when compared to DMPP, point towards a general shut-down of the cell by ATU, possibly towards a state of intense quiescence approaching dormancy^54,55^. A surprising feature in this state of the cell is the up-regulation of the translation initiation factor eIF-2B. This is intriguing as no other translation initiation factors exhibit this response, which is counter to the coordinated response of these same proteins in a recent carbon limitation study of *N. viennensis*^27^. Additionally, this is only observed at the protein level as the corresponding transcripts show no differential expression. This lone response could be tied to the complicated evolutionary history of eIF-2B in archaea in which this protein can also interact with sugars to detect the energy status of the cell^56^. While intriguing, interpretation must be done cautiously as this protein was below the detection limit in DMPP and control cultures and its exact fold change can only be estimated (see supplementary materials and methods). Nevertheless, the metabolic transition to a state resembling dormancy or quiescence is supported by the inability of the cells to recover when additional ammonia or urea is added. Unlike with DMPP, this suggests a form of non-competitive inhibition in which ATU disables the AMO through a means other than physically blocking ammonia from reaching the active site, as previously suggested for AOB^57,58^.

### Models for inhibition and cellular responses for DMPP and ATU

These combined observations allowed for the development of a detailed model of inhibition for DMPP and ATU that was not previously possible. DMPP is proposed to physically block ammonia from reaching the active site of the AMO complex, but not covalently bind, thereby initiating a response of the cell that is akin to low substrate availability (Fig. 8). The cell then invests in alternate mechanisms for obtaining ammonia (i.e. urease and urea transporters) while fortifying its existing machinery through the up-regulation of additional *amoC* subunits and the expression of multiple chaperone heat shock proteins. Conversely, ATU inhibits the AMO through an alternative mechanism that cannot be relieved with additional substrate. A likely possibility is the binding of the thiocarbonyl group of ATU with the copper centers found in the AMO complex. This would be particularly effective if the copper in the archaeal AMO Cu_C_ and Cu_D_ copper sites is found in its reduced form (Cu(I)) as hypothesized for the homologous AMO of bacteria^59^. A mechanism such as this would not be responsive to additional ammonia, as observed here. This type of interaction that leads to an inactive AMO likely initiates a cellular response towards quiescence by an unknown molecular mechanism.

**Figure 8.**
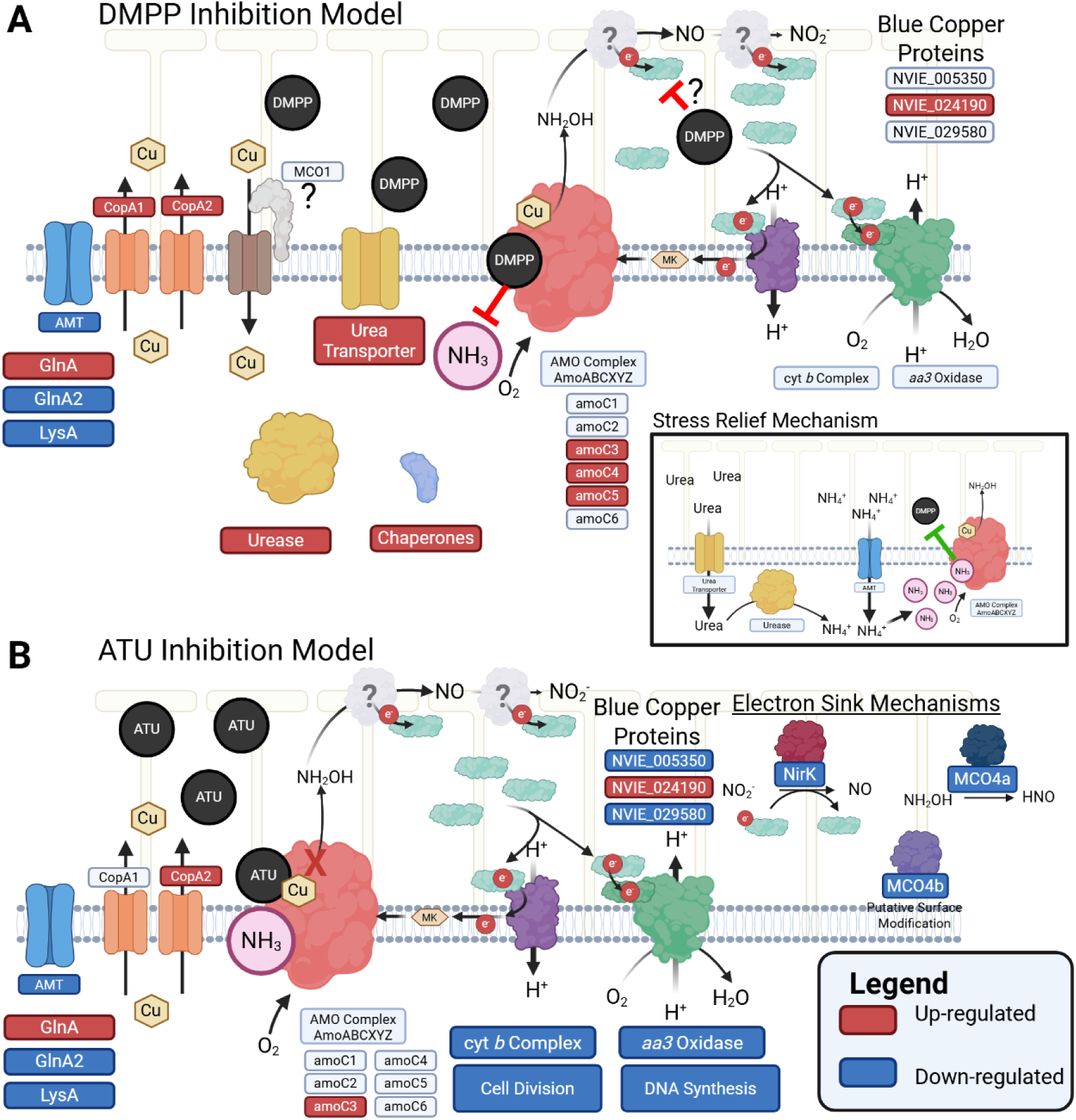
Model of proposed inhibition mechanisms for DMPP (A) and ATU (B) modes of action. **A)** A model of competitive inhibition is proposed for DMPP supported by the up-regulation of urease and urea transporters and the relief of stress by added ammonium or urea. **B)** A model of non-competitive inhibition is proposed for ATU that causes a more severe physiological response supported by the down-regulation of many electron transfer chain genes and proteins and the inability to relieve stress through added copper, metals, ammonium, or urea. Locus tags and accession numbers for all genes and proteins can be found in Dataset S1. Created in BioRender. Hodgskiss, L. (2026) https://BioRender.com/tsocl6b.

## Conclusion

This study used a combination of functional omics and physiological tests to clarify the effects of the NIs DMPP and ATU on the ammonia-oxidizing archaeon *N. viennensis*. To the authors’ knowledge, this represents the first time the physiological effects of NIs have been investigated at a functional omics level of any nitrifier. The results presented here overturn the assumptions that the primary mechanism of DMPP and ATU is the chelation of free copper and propose a method of competitive inhibition modeled for DMPP and non-competitive inhibition represented by ATU in AOA. Both modes of inhibition were associated with clear transcriptomic and proteomic signals that will be informative for the evaluation of future synthetic and biological NIs. While the concentration of DMPP and ATU applied to *N. viennensis* are high in comparison to common practice, the physiological information gained from this study is insightful. In particular, the demonstration that the effectiveness of NI stress can potentially be influenced by high substrate levels is important to take into account when looking for future novel NIs that would be used together with high amount of fertilizer. Furthermore, the up-regulation of urease vs. the down-regulation of the terminal oxidase can serve as initial indicators of the possible mechanisms of newly identified nitrification inhibitors for AOA.

## Materials and Methods

### Strain, growth conditions and nitrification inhibitor preparation

*Nitrososphaera viennensis* EN76, isolated from a garden soil in Vienna, Austria^21^, was used for all experiments. *N. viennensis* EN76 was grown aerobically in liquid batch culture, in the dark with shaking (80 rpm), at an optimum temperature of 42°C and pH 7.5, in 10 mM HEPES-buffered fresh water medium (FWM), supplemented with 2 mM NH_4_Cl. *N. viennensis* growth was monitored via nitrite measurements as previously described^29^. DMPP and ATU were dissolved in autoclaved sterile ddH_2_O before being added in the necessary volume to achieve the target concentration for the respective experiments. Stock solutions for the nitrification inhibitors (NIs) were freshly prepared before each experiment.

### Micro-respiration assays

To determine whether the two NIs are targeting the first or the later steps of nitrification, a micro-respiration assay (see details in supplementary materials and methods) was conducted with 50x concentrated cultures of *N. viennensis* in FWM medium, either supplemented with 200 µM NH_4_Cl or 200 µM NH_2_OH as substrate and the addition of 5 mM of DMPP or 3 mM of ATU to achieve a strong or complete inhibitory effect (see supplementary materials and methods). Concentrations were determined based on previous work with DMPP^60^ and ATU^30,38^ in archaeal ammonia-oxidizers. Upon addition of substrates and inhibitors, oxygen consumption was monitored using a microoptode (Unisense, Denmark) and used as an indicator of metabolic activity.

### Time-series response of *N. viennensis* to DMPP

A time-series experiment was carried out with DMPP to determine the response time needed to observe changes in both transcripts and proteins in *N. viennensis*. This compound was chosen as it is a well-studied NI with previously established EC_50_ values in neutrophilic AOA cultures. Cultures of *N. viennensis* were grown (250 mL culture in 500 mL Schott bottles) under the standard conditions described above and exposed to 2 mM DMPP. Triplicate cultures treated with DMPP were harvested by filtration at 0.5, 1, 3, 6, and 12 hours post application of DMPP and compared to control cultures harvested at 0.5 h. A single control at the beginning of the experiment was considered sufficient due to the short duration of the experiment in relation to the organisms doubling time (∼14 hours). Two standard cultures (no DMPP) and two treated with DMPP were not harvested and used for monitoring AOA growth via nitrite measurements. Filtered biomass was stored frozen at -70 °C until the simultaneous extraction of RNA and proteins was performed (supplementary materials and methods).

### Transcriptomic and proteomic studies of *N. viennensis* to DMPP and ATU stress

To clarify the physiological effect of two different NIs on an AOA, cultures of *N. viennensis* were grown under the same conditions as described above (250 mL culture in 500 mL Schott bottles) and exposed to DMPP and ATU for proteomic and transcriptomic comparative analysis. Eight replicates were grown for each condition (control, DMPP and ATU), five of which were used for omics analysis and three to monitor growth via nitrite measurements. The NIs were applied at a final concentration of 2 mM for DMPP and 0.5 mM for ATU to achieve similar but not complete growth impairment at the mid-exponential growth phase. The concentration of DMPP was chosen from the half-maximal effective concentration (EC_50_) of the soil AOA *Nitrosocosmicus franklandianus*^60^ and the concentration of ATU was estimated based on previous work with *N. viennensis*^30^. Cultures were harvested at 9 h, i.e. at 70% of *N. viennensis* doubling time, based on the results from the time-series experiment. Biomass was harvested by filtration and frozen at -70 °C before subsequent RNA and protein extraction. Following protein extraction, samples were processed and analyzed by mass spectrometry. The resulting proteomic data were analyzed and interpreted (see supplementary materials and methods).

### Relief experiments

Following the identification of differentially expressed genes and proteins, liquid batch cultures with *N. viennensis* (20 mL cultures in 30 mL Greiner plastic tubes) were exposed to 2 mM DMPP and 0.5 mM ATU in combination with the addition of: copper (up to 3 µM), trace elements (up to 10× the initial concentration), ammonium (5 mM), and urea (up to 2.5 mM), to test recovery from the stress. All conditions were performed in triplicate cultures.

## Data Availability

Transcriptomic data for the time-series experiment and comparison of DMPP and ATU have been deposited in the NCBI Sequencing Read Archive (SRA) under the BioProject numbers PRJNA1450353 and PRJNA1450355, respectively. The mass spectrometry proteomics data have been deposited to the ProteomeXchange Consortium^61^ via the PRIDE^62^ partner repository with the dataset identifiers PXD077624 and 10.6019/PXD077624 for the DMPP time-series and PXD077699 and 10.6019/PXD077699 for the DMPP and ATU comparison. Raw data for transcriptomics (input and output of DESeq2) and proteomics (output of MaxQuant and processed data) can be found in Dataset S4.

## Acknowledgements

We thank Assistant Prof. Vasileiadis Sotirios of the University of Thessaly for technical assistance and Dr. Eleftheria Bachtsevani for laboratory support. The RNA sequencing was performed by the Next Generation Sequencing Facility at Vienna BioCenter Core Facilities (VBCF), member of the Vienna BioCenter (VBC), Austria. The computational results of this work have been achieved using the Life Science Compute Cluster (LiSC) of the University of Vienna.

