## Supplementary Material for "Common nitrification inhibitors exhibit varied physiological mechanisms on an ammonia-oxidizing microorganism"

#### Contents:

Supplementary Results and Discussion

Supplementary Materials and Methods

Supplementary Figures

References

**Additional datasets:**

Dataset S1-Detailed Growth, Gene, and Protein Data from Main Figures

Dataset S2-Transcriptomic Datasets

Dataset S3-Proteomic Datasets

Dataset S4-Raw Data

### Supplementary Results and Discussion

**Response of genes and proteins in DMPP time-series experiment**

The strong inhibition of growth by the exposure of *Nitrososphaera viennensis* to 2 mM DMPP prompted an examination of the response of energetic complexes within the time-series experiment. This analysis revealed a broadly consistent response pattern of down-regulation, with the notable exception of the ammonia monooxygenase (AMO) complex (Fig. S11). Most AMO subunits remained at transcript levels comparable to controls, except for the auxiliary *amoC3* and the primary *amoC6* gene[1], which increased over time. Other energetic complexes exhibited rapid down-regulation within the first 30 min. Complex I decreased initially and then stabilized without further significant changes and was found in clusters 1 and 2. Complexes II (clusters 1 and 2), III (cluster 2 and 3, although the line plots indicate a clearly decreasing trend throughout the time series), and IV (cluster 2 and *coxA2* in cluster 3) showed immediate down-regulation followed by partial recovery at the final time point, although this recovery was not statistically significant for all genes. In contrast, Complex V showed a continuous decline throughout the experiment and was found exclusively in cluster 1 (Fig. S12).

To further identify genes and proteins of potential interest, clusters with comparable patterns in transcriptomic and proteomic data were compared. Specifically, clusters with increasing or decreasing relative abundance in both RNA and proteins were cross-examined to point out common patterns for genes and their respective products. Common decreasing RNA/proteins included ribosomal proteins (*rps3p* and *rpl2p*), threonine synthase (*thrC2*), *leuC2* and a DEAD box RNA helicase (NVIE_022610). Genes and proteins with transient induction included glutamine synthetase (*glnA*), an SNF7 domain containing protein (NVIE_018700, *cdvB*-3), a putative protease from the DO family protein (NVIE_022330) and a methyltransferase type 12 (NVIE_007040). The identification of *glnA* at both the RNA and protein level prompted an additional look at the alternative gene *glnA2*. At the RNA level *glnA2* decreased, the opposite response observed for *glnA* (Fig. S5). At the protein level, however, GlnA2 remained relatively constant across the time-series.

A look at genes and proteins that both increased revealed six genes with concordant increasing trends at both levels: *ureC1* (urease subunit alpha), *flaB3* (archaeal flagellin), *cheB* (chemotaxis response regulator protein – glutamate methylesterase), NVIE_010090 (dehydrogenase – close to chemotaxis and flagella clusters), NVIE_009790 (putative methyl-accepting chemotaxis sensory transducer), and NVIE_004270 (hypothetical protein). The identification of *ureC1*, the primary subunit of urease, amongst the genes showing a clear progressive upregulation pattern, prompted a closer look at other urease genes. At the transcript level, the primary urease operon genes (*ureC1*, *ureB1*, and *ureA1*) and the accessory gene *ureG* all responded rapidly to DMPP exposure and were found within cluster 3 (*ureA1B1C1*) and cluster 6 (*ureG*), with changes detectable within 0.5 hours and a continuous increase in expression up to 12 h (Fig. S13). At the proteomic level, only *ureC1* was statistically significant within the ANOVA analysis. However, *ureB1* and *ureG* were still detected in all proteomic data, and although their abundance followed a similar increasing trend, protein-level changes became apparent only after 3 h of exposure, indicating a delayed translational response of 3-6 hours, similar to the broader observation found in the PCA plots (Fig. 2C).

A broader look at motility genes was also warranted based on the high-enrichment of this arCOG category in cluster 3. This arCOG category pointed towards a specific motility cluster within the *N. viennensis* genome: NVIE_009600-NVIE_010090. These motility-related genes (including flagellar, chemotaxis, and neighboring genes) generally displayed increasing transcript abundance, particularly after 1 h of DMPP exposure. Notably, four genes (*cheB*, *flaB3*, NVIE_009790, and NVIE_010090) showed consistent trends at both RNA and protein levels (Fig. S13), with transcriptional up-regulation preceding detectable increases in protein abundance by at least 3 hours. While motility genes demonstrated a strong up-regulation after exposure to DMPP in the time-series experiment, this trend was not observed when in the later comparative experiment performed with ATU. Surprisingly, none of the motility genes identified in the time-series showed any differential expression, and two of the corresponding proteins were down-regulated in the proteome when compared to controls (CheB and NVIE_009790). Further investigation of the controls of each experiment revealed a much higher general expression of this motility cluster in the time-series experiment, whereas the motility cluster in the later comparative experiment was much lower (Fig. S14). One possibility for these differences could be the time at which each experiment was sampled in the growth curve. The time series experiment was harvested when cultures had reached approximately 500-700 µM nitrite while the comparative experiment was started at a slightly higher level of approximately above 1100 µM nitrite.

Due to these differences, motility was not considered to be a result of DMPP stress. Unlike the motility gene cluster, other responses to DMPP were highly conserved in both experiments (i.e., urease, urease accessory genes, transporters). Therefore, these physiological effects were taken to be a specific response to DMPP rather than the growth phase. However, direct comparisons between the time-series and comparative experiment should be treated cautiously as the time-series was not designed for this purpose.

### Supplementary Materials and Methods

**_Nitrososphaera viennensis_ EN76, growth conditions and chemicals**

*Nitrososphaera viennensis* EN76, isolated from a garden soil in Vienna, Austria[2], was used for all experiments. *N. viennensis* EN76 was grown aerobically in the dark with shaking (80 rpm), at an optimum temperature of 42°C and pH 7.5, in 10 mM HEPES-buffered fresh water medium (FWM) supplemented with 2 mM $NH_4^+$ ($NH_4Cl$). FWM consisted of NaCl (1 g $L^{-1}$), $MgCl_2 \cdot 6H_2O$ (0.4 g $L^{-1}$), $CaCl_2 \cdot 2H_2O$ (0.1 g$\cdot L^{-1}$), $KH_2PO_4$ (0.2 g $L^{-1}$) and KCl (0.5 g $L^{-1}$). Modified non-chelated trace element mixture consisted of HCl (100 mM), $H_3BO_3$ (0.03 g $L^{-1}$), $MnCl_2 \cdot 4H_2O$ (0.1 g $L^{-1}$), $CoCl_2 \cdot 6H_2O$ (0.19 g $L^{-1}$), $NiCl_2 \cdot 6H_2O$ (0.024 g $L^{-1}$), $CuCl_2 \cdot 2H_2O$ (0.002 g $L^{-1}$), $ZnSO_4 \cdot 7H_2O$ (0.144 g $L^{-1}$) and $Na_2MoO_4 \cdot 2H_2O$ (0.036 g $L^{-1}$). After autoclaving FWM, final medium was prepared with the addition of non-chelated trace element mixture (0.1% dilution), FeNaEDTA solution (7.5 µM), ammonium chloride (2 mM), sodium bicarbonate (2 mM) and pyruvate (1 mM) under sterile conditions. The pH of the medium was 7.5 with the addition of HEPES buffer (1M HEPES, 0.6M NaOH) to a final concentration of 10 mM. The absence of any contamination was verified by adding 50 µL of the grown cultures onto R2A agar plates and incubating them under the same conditions as the cultures[3]. The nitrification inhibitors (NIs) used were 3,4-dimethylpyrazole phosphate (DMPP, Cayman Chemical Company, purity ≥ 98%) and allylthiourea (ATU, Sigma-Aldrich, purity 98%). DMPP and ATU were dissolved in autoclaved sterile dd$H_2O$ before being added in the necessary volume to achieve the target concentration. Stock solutions for the NIs were prepared fresh before each experiment. The total volumes of NIs added to the cultures were not more than 0.1% of the culture volume and were therefore considered to be a negligible volume addition.

**Nitrite measurements**

Nitrite concentrations were determined colorimetrically at 540 nm in a 96-well plate format assay by diazotizing and coupling with Griess reagent as previously described[4]. For each measurement, standards of the appropriate curves of $NaNO_2$ were used.

**Micro-Respiration Assay**

A high-precision Micro-Respiration system (Unisense, Denmark) fitted with a microoptode was employed to monitor oxygen levels, enabling the determination of oxygen consumption rates (respiration) in small, sealed systems. Custom-built chambers, approximately 15 mm in diameter and 2 mL in volume, were used. Each chamber was sealed with a lid containing a capillary aperture of roughly 0.7 mm × 10 mm. Because the chambers were handcrafted, their exact volumes were measured individually. During measurements, the chambers were placed in an MR2-Rack equipped with a magnetic stirrer. The microsensor was mounted in an aluminum sensor guide with adjustable plastic tips. This guide was positioned between the chamber rack and the MR-lid, allowing the sensor tip to be precisely aligned and inserted through the capillary opening into the sealed chamber. Before the experiments, the oxygen microsensor was calibrated using a two-point approach. The zero-point was set using an anoxic solution prepared with sodium ascorbate and NaOH, both to final concentrations of 100 µM, while the 100 % air-saturation calibration was performed in a separate chamber containing FWM medium, continuously aerated with an air bubble generator and maintained at 42 °C in a water bath. For the experiments, the respiration chambers were loaded with 50× concentrated late exponential phase cultures of *N. viennensis* in FWM medium, either supplemented with 200 µM $NH_4Cl$ or 200 µM $NH_2OH$. In tests with DMPP, HEPES concentration was increased from 10 mM to 50 mM to prevent a pH drop due to the high amounts of DMPP. Fifty-times concentrated late exponential phase cultures were chosen based on previous tests for the optimal cell density. This concentration showed the highest nitrite production while keeping the incubation time low to prevent the bias of $NH_2OH$ reactivity. Cell abundance was analyzed by flow cytometry as explained in detail in Malits et al.[5]. Oxygen consumption was monitored with the microoptode while the chamber contents were stirred at 600 rpm and kept at a constant 42 °C using a water bath. To avoid light interference with the sensor, the water bath was kept in darkness. DMPP (5 mM) or ATU (3 mM) was added to the chambers in combination with either 200 µM $NH_4Cl$ or 200 µM $NH_2OH$. Higher final concentrations of each NI were used to account for the hich cell density of the cells, as previous experiments were done in liquid batch cultures and not in concentrated cells.

Following initial experiments, relief of stress by extra ammonium was tested by the addition of 7 mM to chambers containing 200 µM of ammonium and either 5 mM of DMPP or 3 mM ATU and compared against a control culture with 7 mM of ammonium with no added inhibitor.

DMPP was additionally tested at concentrations of 7 mM with both 200 µM of ammonium and 200 µM of hydroxylamine to achieve higher inhibition. Relief at these concentrations was also tested with the addition of 14 mM ammonium (ammonium test) and 5 µM of copper (ammonium and hydroxylamine tests). All relief experiments were compared to the appropriate controls.

##### Stress relief experiments in liquid batch cultures

Cultures of *N. viennensis* were grown in triplicates (0.25% inoculum), in 20 mL of medium in leak-resistant Thermo Scientific™ Sterilin™ Quickstart Universal Polystyrene 30mL containers and exposed to DMPP (2 mM) and ATU (0.5 mM) in the presence of different concentrations of copper, trace elements, ammonium and urea, to assess whether the stress caused by the NIs can be relieved. NIs (2 mM of DMPP or 0.5 mM of ATU) were added at early exponential phase (120 to 200 µM $NO_2^-$). Cultures serving as controls were amended with urea, ammonium, or copper at the same time that NIs were added. Amendments to cultures treated with NIs were added two days after exposure to DMPP or ATU. Two different sources of copper (copper sulfate or copper chloride, 3000 nM) were used to ensure that any possible effect would be caused by copper and not from the accompanying anion. Trace elements were added at 10x the initial concentration to test recovery from other metals. 5 mM of ammonium and 1 or 2.5 mM urea (corresponding to 2 and 5 mM ammonium upon hydrolysis, respectively) was added to test whether there would be recovery with extra substrate. In the NI-treated cultures, all the solutions were added 2 days after incubation with the NIs. Nitrite was measured daily until recovery or stationary phase. In cultures receiving 2.5 mM of urea, ammonium concentration was measured at day 11 for the control and day 9 for the NIs after urea addition to ensure conversion to ammonium.

**Liquid batch culture bioassay for recovery experiments**

Triplicate cultures of control, DMPP and ATU conditions, at the same concentrations as the stress relief experiments, were prepared as described above. One day after the start of the incubation with each NI, 2 mL of culture was concentrated by centrifugation (15,000 x g, 20 minutes, 4°C). The pellet was washed twice with medium without substrate and trace elements. Washed pellets were resuspended in 1 mL of medium. The inoculation volume of new cultures was normalized based on the lowest average nitrite level to achieve similar starting cell densities. Each condition was inoculated in fresh medium with the corresponding amendments and nitrite was monitored.

**Time-Series Experiment with DMPP**

Cultures of *N. viennensis* were grown (0.25% inoculum) in 500 mL DURAN® Original GL 45 laboratory bottles, with screw cap and pouring ring and polypropylene cap containing 250 mL of culture and exposed to DMPP for proteomic and transcriptomic analysis.

DMPP was added at a final concentration of 2 mM[6] when cultures had produced approximately 500 µM nitrite to obtain sufficient biomass for RNA and protein extraction. For the assay, control cultures without DMPP were included. The cultures treated with the NI were harvested by filtration at 0.5, 1, 3, 6, and 12 hours post application of DMPP and compared to a control culture harvested at the same time as the 0.5 h treated culture. The control time point was selected to serve as a baseline reference for the subsequent time points to determine transcriptional and proteomic changes over time following DMPP exposure. Triplicate cultures were harvested for the control and each time point. Two cultures each for control and DMPP conditions were not harvested to serve as growth controls. Cells were collected by filtration onto MCE Membrane Filter (0.2 µm pores and 45 mm diameter) using a vacuum pressure bottle top apparatus and subsequently used for RNA and protein extraction. The filters with the attached biomass were placed in 50 mL falcon tubes and immediately transferred to -70°C until further handling.

**Growth with DMPP and ATU for Comparative Omics**

Following the identification of the optimal time of harvesting to capture the effects in both transcriptome and proteome from the same sample following NI addition, cultures of *N. viennensis* were grown in 8 replicates (0.25% inoculum) and exposed to 2 mM of DMPP[6] and 0.5 mM of ATU[7]. Five replicates were used for omics and three were used to monitor the growth and inhibition of *N. viennensis*. The NIs were applied at a final concentration of 2 mM for DMPP and 0.5 mM for ATU, when cultures had produced approximately 700 µM nitrite to obtain sufficient biomass for RNA and protein extraction. Control cultures with no NIs were grown in quintuplicates. Cells were harvested by filtration and filters were stored in 50 mL falcon tubes and frozen at -70 °C after 9 hours until the extraction of RNA and proteins.

#### Dual extraction of RNA and proteins

RNA was extracted using the mirVana™ miRNA Isolation Kit. The MCE filters with frozen biomass were washed and resuspended thoroughly in 700 µL of Lysis/Binding Buffer using a 1 ml syringe with a needle for 5 minutes per sample in the 50 mL falcon used for storage and then placed on ice for 10 minutes incubation. The mixture was transferred into RNAse/DNAse-free 2 mL Eppendorf tubes, 60 µL of Homogenate Additive was added, vortexed thoroughly and then incubated for 10 minutes on ice. 600 µL of acid phenol-chloroform was added and then vortexed for 60 seconds (from this point on the entire protocol was performed in a fume hood). Samples were centrifuged at 10,000 x g for 5 minutes at room temperature. The upper (aqueous) phase, containing RNA, was carefully transferred, without touching the intermediate or lower phase, to a new 2 mL Eppendorf tube. At this point the lower, non-aqueous phase, containing the protein fraction, was stored on ice until RNA extraction was completed. To continue the isolation of RNA, 700 µL of 100% ethanol at room temperature was added to the aqueous phase and mixed thoroughly by vortex for 30 seconds. The mixture was passed through a filter into a cartridge supplied with the kit and centrifuged at 10,000 x g for 15 seconds. The flow-through was discarded (this step was repeated until no more mixture was left). The filter was washed with 700 µL of miRNA Wash Solution 1, centrifuged for 15 seconds at 10,000 x g and the flow-through discarded. The filter was then washed with 500 µL of miRNA Wash Solution 2/3, centrifuged for 15 seconds at 10,000 x g and the flow-through discarded.  The wash step with Solution 2/3 was repeated. The filter was placed into a new sterile RNAse/DNAse-free Eppendorf and 60 µL of preheated 95°C DEPC dd$H_2O$ was added. Samples were incubated for 10 minutes at room temperature and then centrifuged at 10,000 x g for 1 minute. The RNA was left to homogenize for 30 minutes on ice before measuring the concentration and possible phenol contamination using a NanoDrop (NanoPhotometer N60, Implen).

Following the RNA extraction, the protein extraction was continued with the non-aqueous phase. 1.5 mL of 0.1 M ammonium acetate in methanol with 0.5% β-mercaptoethanol was added to the samples, the tubes were inverted up and down to gently mix the solution and were then stored overnight at -20°C to precipitate proteins.

**Phenol Cleanup of the RNA samples**

If phenol peaks were observed when measuring the RNA concentration, the samples were cleaned following a specific procedure. 240 µL of DEPC dd$H_2O$ was added to the samples, to a total volume of 300 µL, including 20 µL of sodium acetate (3M) and 1 µL of glycogen (RNA grade), followed by mixing by vortexing. Then 550 µL of ice-cold 100% ethanol was added, vortexed and incubated at -70 °C for 30 minutes. Mixtures were centrifuged at 11,000 x g for 20 minutes at 4°C to precipitate RNA. The supernatant was discarded. Residual liquid was carefully removed and discarded using a 10 µL pipette and samples were left to dry for 10 minutes. Afterwards RNA was resuspended in 60 µL of DEPC dd$H_2O$ and the concentration was re-measured. Samples were frozen at -70 °C until further processing by DNAse (see below).

**DNA digestion for the RNA samples**

All RNA samples were treated for DNA digestion. 6.8 µL of Turbo DNAse buffer and 1-2 µL of Turbo DNAse were added to 60 µL of RNA solution. For 10 µg of RNA, 1 µL (2 U) was used; if RNA amount exceeded 10 µg, 2 µL was used. Samples were incubated at 37 °C for 2 hours to ensure complete DNA digestion. Afterwards, 6.8 µL of Inactivation Reagent was added, tubes were gently mixed several times and centrifuged at room temperature for 1.5 minutes at 11,000 x g. The upper, RNA-containing phase was transferred into a new DNAse/RNAse-free 1.5 mL Eppendorf tube. The presence of remaining DNA was checked by PCR. PCR was performed using AOA-specific primers targeting the gene NVIE_000940 (cell surface protein with DUF11 domain) with the following primer set:

forward primer 5'-CGCGTCTGCCGTGATTATTG-3', reverse primer 5'-ATAGACCTGTCTAGCGGCCA-3'. A total of 1 µL of sample was tested in a 20 µL reaction volume along with the following: 0.4 µL of 10 mM dNTP mix (Thermo Fisher), 0.2 µL of Phusion™ High-Fidelity DNA Polymerase (NEB), 4 µL of 5X Phusion GC Buffer, 0.5 µL of forward primer stock (10 µM stock concentration), 0.5 µL of reverse primer stock (10 µM stock concentration), and DNA-free water. The following PCR conditions were used: 95 °C for 5 min followed by 30 cycles of 95 °C for 30 s, 58 °C for 30 s and 72 °C for 20 s, and a final elongation step at 72 °C for 10 min. Samples were then analyzed by agarose gel electrophoresis (1%, 240 V for 45 minutes) to verify complete digestion of DNA in the samples.

**Protein washing**

After incubation overnight at -20 °C, samples were centrifuged for 15 minutes at 4 °C, at maximum speed using a benchtop centrifuge. The supernatant was discarded and 1.8 mL ice-cold methanol was added. Samples were placed in a cooled ultrasonic water bath (Transsonic 700/H water bath (Elma) sonicator) until the pellet was fully resuspended. Ice was placed in the ultrasonic bath to prevent overheating, and samples were checked every 4 to 5 minutes to ensure complete resuspension; if necessary, samples were gently mixed. The samples were then centrifuged for 10 minutes at 4 °C, at maximum speed and the supernatant was discarded. Methanol wash was repeated once more. After discarding the supernatant, 1.8 mL of ice-cold acetone was added and samples were sonicated again. After sonication samples were centrifuged for 15 minutes at 4°C, at maximum speed. Supernatant was discarded and the resulting pellet was air-dried under a fume hood for approximately 10 minutes (until pellets were dry). Samples were stored at -70 °C until quantification and digestion.

**Time-Series Experiment Protein Quantification and Peptide Digestion**

Protein concentration was quantified with the "Pierce® Microplate BCA Protein Assay Kit – Reducing Agent Compatible, followed by digestion using the PreOmics iST kit. To quantify proteins, 50 µL of LYSE buffer from the PreOmics kit was added to the washed pellet for the samples to be quantified. After determining the concentration, the amount needed for 25 µg of protein was transferred to an Eppendorf tube and more LYSE buffer was added to a final volume of 50 µL. Samples were then placed in a heating block at 95 °C, 1000 rpm for 10 minutes. The sample was then transferred to a filter cartridge supplied by the PreOmics kit.

To prepare the digestive enzymes, 210 µL of RESUSPEND buffer was added to DIGEST and was shaken at 500 rpm, room temperature for 10 minutes. 50 µL of the prepared DIGEST solution was added to each sample in the filter cartridges and placed in a pre-heated heating block at 37 °C, 500 rpm for 3 hours. Following this, 100 µL of STOP buffer was added to each sample and samples were shaken at 500 rpm, room temperature for 1 minute. If necessary, samples were stored overnight at -20 °C before the washing steps performed in the following day.

Samples were centrifuged at 3,800 rcf for 1-3 minutes until complete flowthrough of solution. For peptide washing, 200 µL WASH buffer 1 was added to the filter cartridge and centrifuged at 3,800 rcf for 1-3minutes. Following this, 200 µL WASH buffer 2 was added to the filter cartridge and centrifuged at 3,800 rcf for 1-3minutes. For elution of peptides, 100 µL of ELUTE buffer was added to the cartridge and transferred to a new Eppendorf tube and centrifuged at 3,800 rcf until complete flow-through. The elution step was repeated with 100 µL of ELUTE buffer and collected within the same tube resulting in a final volume of 200 µL with eluted peptides. Samples were dried in a speedvac until complete dryness of the digested peptides. Digested samples were stored at -70 °C until mass spectrometry analysis.

#### DMPP and ATU Comparison Experiment Protein Quantification and Peptide Digestion

Protein quantification was performed using the “Pierce® Microplate BCA Protein Assay Kit – Reducing Agent Compatible as described above. The PreOmics iST protocol was modified with guidance from the company for improved results and removal of mass spectrometry contaminants as follows (changes in bold):

After determining the concentration, the amount needed for 25 µg of protein was transferred to an Eppendorf tube and more LYSE buffer was added to a final volume of 50 µL. Samples were then placed in a heating block at 95 °C, 1000 rpm for 10 minutes.

To prepare the digestive enzymes, 210 µL of RESUSPEND buffer was added to DIGEST and was shaken at 500 rpm, room temperature for 10 minutes. **50 µL of the prepared DIGEST solution was added to each sample in a protein LoBind Eppendorf tube** and placed in a pre-heated heating block at 37 °C, 500 rpm for 3 hours. Following this, 100 µL of STOP buffer was added to each sample and samples were shaken at 500 rpm, room temperature for 1 minute. If necessary, samples were stored overnight at -20 °C before the washing steps performed in the following day.

Samples were centrifuged at 3,800 rcf for 1-3 minutes until complete flowthrough of the solution. For peptide washing, 200 µL WASH buffer 1 was added to the filter cartridge and centrifuged at 3,800 rcf for 1-3minutes. Following this, 200 µL WASH buffer 2 was added to the filter cartridge and centrifuged at 3,800 rcf for 1-3 minutes**. Peptides were washed for a second time with WASH buffer 2.** For elution of peptides, 100 µL of ELUTE buffer was added to the cartridge and transferred to a new Eppendorf tube **and centrifuged at 1000 rcf** until complete flow-through. The elution step was repeated with 100 µL of ELUTE buffer and collected within the same tube resulting in a final volume of 200 µL of eluted peptides. **Eluted samples were then centrifuged for 10 minutes at**

**2,250 rcf to precipitate any remaining mass spectrometry contaminants. The supernatant (190-195 µL) was transferred to a new protein LoBind Eppendorf tube.** Samples were dried in a speedvac until complete dryness of the digested peptides. Digested samples were stored at -70 °C until mass spectrometry analysis.

#### Liquid chromatography–mass spectrometry (LC–MS)

Purified tryptic peptides (1 µg) were analyzed by LC–MS using a Thermo Scientific Ultimate 3000 nano LC system for reversed-phase separation, coupled to a Q Exactive Plus Orbitrap Mass Spectrometer (Thermo Fisher Scientific). Peptides were separated on an Easy-Spray PepMap RSLC C18 Column (2 µm, 100 Å, 75 µm × 50 cm) at 55 °C and 300 nL $min^{-1}$ using a 5–35% mobile phase B gradient over 105 min, followed by 80% mobile phase B over 5 min (mobile phase A: 0.1% formic acid in water; mobile phase B: 89.9% acetonitrile, 0.1% formic acid, 10% water). MS data were collected in positive ion mode using top-20 data-dependent acquisition. Full MS scans were acquired at 70,000 resolution (AGC $1 \times 10^6$, 120 ms, m/z 350–1600), and Tandem MS spectra were acquired at 17,500 resolution (AGC $5 \times 10^4$, 1.6 m/z isolation window, and 100 ms) using HCD (NCE 27%). Dynamic exclusion was set to 60 s.

#### Transcriptomic analysis

RNA was submitted to the Vienna Biocenter Facility (VBC) for NovaSeq S4 PE150 XP (Illumina) sequencing with a species-specific rRNA depletion step. Generated raw reads were downloaded from VBC, checked using md5sum, and were processed and mapped to the available microbial genome: *Nitrososphaera viennensis* EN76, genome assembly ASM69878v1 using the GenBank annotation files. FastQC v.0.12.1[8] and multiQC v.1.25[9] were used to initially analyze the data. Reads were trimmed twice using fastp v.0.23.4[10] to fully remove adapter sequences. The following specific settings were used for the first trimming: --average_qual 30 --detect_adapter_for_pe --trim_poly_g, and the following for the second trimming: --average_qual 30  --length_required 30 --trim_front1 12 –trim_front2 12. rRNA reads were sorted out with SortmeRNA[11] with following specific settings: --fastx –aligned –other –paired_out –out2, and remaining RNA reads were mapped and counted to the specific genome using Salmon 1.10.3[12]. For index creation in Salmon, a transcript file was created using gffread using default settings with the GFF (annotation file) and FNA (genome file_ as inputs.  The transcript and FNA file were then concatenated (Nvie_gentrome.fa) and used as input when creating a decoy-aware index file for Salmon using the following commands: salmon index -t Nvie_gentrome.fa -d decoys.txt -i Nvie_with_decoys_index –-type puff -k 31, where decoy.txt contains the headers from the FNA file. After index creation, the following specific settings for the mapping and counting: salmon quant -i ./ Nvie_with_decoys_index -l ISR -1 -2 -p 8 –validateMappings –allowDovetail -o. All further analyses were conducted in R 4.5.3[13] using R-Studio version 2025.02.0+496[14].

Raw counts from Salmon were rounded to the nearest integer value were analyzed for differential expression using DESeq2[15] employing the default settings. For multivariate analysis, data was normalized using the rlog function in DESeq2. PCA plots were produced using data from all detected genes from the normalized data using the R packages ggplot2[16], ggrepel[17], and factoextra[18]. Transcripts per million (TPM) values were calculated within the Salmon package. $Log_2$ of TPM was calculated to use for color coding of TPM values when looking at genes individually.

For analysis of the time-series RNA data, a Likelihood Ratio Test (LRT) was performed with the DESeq2 package and the most significant genes (adjusted *P* value < 0.001, Benjamini-Hochberg method) were separated into 6 clusters using Euclidean distances and k-means clustering using rlog TPM counts.  Number of clusters was chosen using an elbow plot method. Results were plotted in a heatmap using ggplot2, factoextra, ggrepel, stringr[19], tidyverse[20], pheatmap[21], NbClust[22], and dplyr[23]. Enrichment analysis of arCOG categories[24] was done using the phyper function to test for enrichment and graphing the results, with the packages tidyverse, magrittr[25], expss[26], RColorBrewer[27], and dplyr. arCOG categories with an adjusted *P* value < 0.05 were labeled as significantly enriched. Venn diagrams were done using Venny 2.1.0 (https://bioinfogp.cnb.csic.es/tools/venny/). Line plots were generated using $log_2$ transformed TPM values. Full time-series LRT results can be found in Dataset S2.

For analysis of the DMPP and ATU comparison, differential expression was calculated based on the default settings of DESeq2. The Benjamini-Hochberg method was used to control for false discovery rate (FDR). Genes with an adjusted *P* value < 0.001 and a log2-FC in treatments of ≥ 1.0 or ≤ -1.0 were considered significantly up/down-regulated. Three univariate comparisons were performed to identify differentially expressed genes: DMPP vs. control, ATU vs. control, and ATU vs. DMPP. Full differential expression results can be found in Dataset S2.

**Proteomic data analysis**

Protein identification from raw mass spectrometry data were performed using MaxQuant version 2.7.5.0[28]. Max missed cleavages was set to 2 with variable modifications of "Oxidation (M)" and "Acetyl (Protein N-term)", fixed modifications set to "Carbamidomethyl (C)", an FDR of 0.01, and the digestive enzyme as Trypsin/P and LysC/P. Minimum number of unique and razor peptides was set to 2 and protein values were LFQ normalized. For Fast LFW settings in the time-series experiment, due to only 3 replicates per condition, LFQ min edges per node was set to 2 and LFQ average edges per node was set to 4. In the comparative DMPP and ATU experiment these Fast LFQ values were left at default settings (LFQ min edges per node was set to 3 and LFQ average edges per node was set to 6). Reference proteome for N. *viennensis* was downloaded from Uniprot in spring of 2019.

For the comparative DMPP and ATU experiment, LFQ data were analyzed using a boxplot analysis for each protein in each condition and the identified outliers were removed and replaced with the corresponding group averages calculated for each protein. Outlier correction was not performed for the time-series data due to the low number of replicates. Afterwards the data were filtered for at least 3 out of 5 replicates with a protein hit in at least one condition for the DMPP and ATU experiment and for at least 2 out of 3 in at least one time point for the time series experiment. The filtered LFQ data was normalized and imputed using the DEP package[29]. Missing data was imputed using the MinProb method with q=0.01 across samples, i.e. missing data was estimated using the bottom 1% of the intensity distribution for each sample. PCA plots of the imputed data were made using the fviz_pca_ind function of the factoextra package in R.

Following imputation, time-series data were analyzed by analysis of variance (one-way ANOVA (aov() function, stats package)) and *P* values were adjusted using the Benjamini-Hochberg method to identify proteins that change over time (adjusted *P* value < 0.05). Data was checked for normality (Shapiro test, shapiro_test() function of rstatix package[30]) and homogeneity of variance (Levene test, leveneTest() function of car package[31]) before being analyzed by ANOVA. Proteins that were found to be significant were separated into 4 clusters using Euclidean distance and k-means clustering. The number of clusters was chosen using an elbow plot method. Line plots were generated using LFQ normalized and imputed values. Full time-series proteomic results can be found in Dataset S3.

For imputed data from the DMPP and ATU comparison experiment, the package DEP was used to perform 3 univariate comparisons: DMPP vs. control, ATU vs. control, and ATU vs. DMPP. Proteins showing a statistical difference were identified using an adjusted *P* value < 0.05 among the tested conditions. Full results, including the number of detected vs. imputed values for each protein, can be found in Dataset S3.

**Sparse partial least square discriminant analysis (sPLS-DA)**

To identify genes or proteins that could potentially distinguish between DMPP and ATU treatments in the DMPP and ATU comparative experiment, a sparse partial least square discriminant analysis (sPLS-DA) was performed. Using the MixOmics[32] package, rlog transformed transcript data or imputed LFQ protein data were analyzed. The analysis was limited to two components and was tuned to identify the smallest number of genes/proteins for each component that would still maximize separation based on the classification error rate. Tuning was validated using the method “Mfold” with folds set to 5, dist set to “max.dist”, and measure set to “overall”. Number of genes/protein was tested from 5 to 200 (RNA) or 5 to 100 (protein) by multiples of 5 with nrepeat set to 100. For RNA, this resulted in 10 genes for variate 1 and 20 genes for variate 2. For protein, this resulted in 5 proteins for variate 1 and 25 proteins for variate 2. Results were used to plot the final sPLS-DA. Results can be found in Dataset S1.

### Supplementary Figures

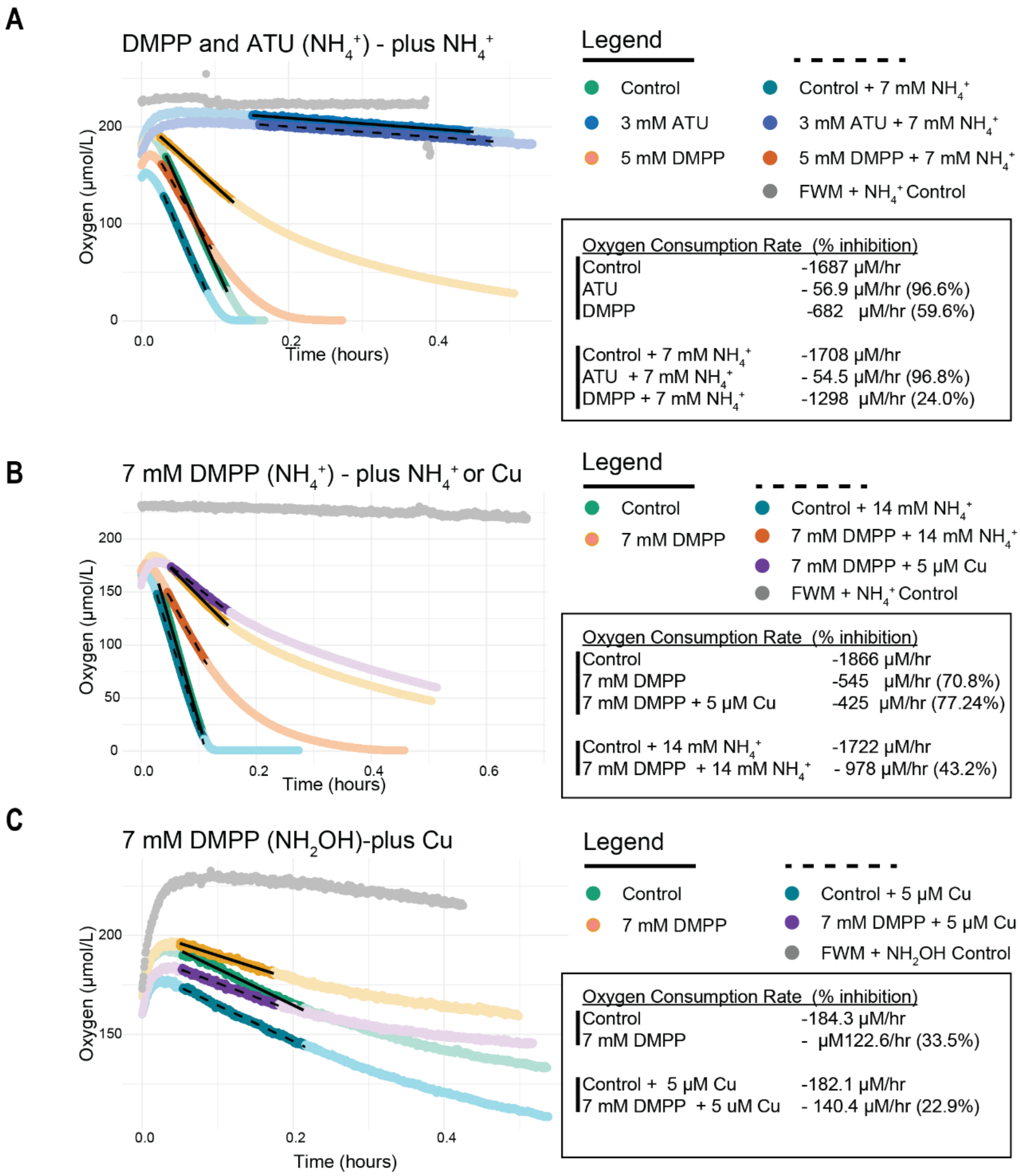


**Figure S1. Micro-respiration assay with stress relief experiments on *N. viennensis*.** **A)** *N. viennensis* exposed to ATU (3 mM) or DMPP (5 mM) in the presence of ammonia (200 µM) as substrate and compared with additions of 7 mM $NH_4^+$. **B)** *N. viennensis* exposed to 7 mM of DMPP and 200 µM of $NH_4^+$ alone or with the addition of 14 mM $NH_4^+$ or 5 µM of $CuCl_2$. **C)** *N. viennensis* exposed to 7 mM of DMPP and 200 µM of $NH_2OH$ alone or with the addition of 5 µM of $CuCl_2$. In all plots, control tests (no additional $NH_4^+$ or Cu) have slopes marked with solid black lines. Relief tests (additional $NH_4^+$ or Cu) are marked with dashed black lines. Percent inhibition was calculated relative to respective controls. Vertical black lines within the "Oxygen Consumption Rate" boxes indicate which samples were compared with which controls. FWM controls represent full medium with no added cells.

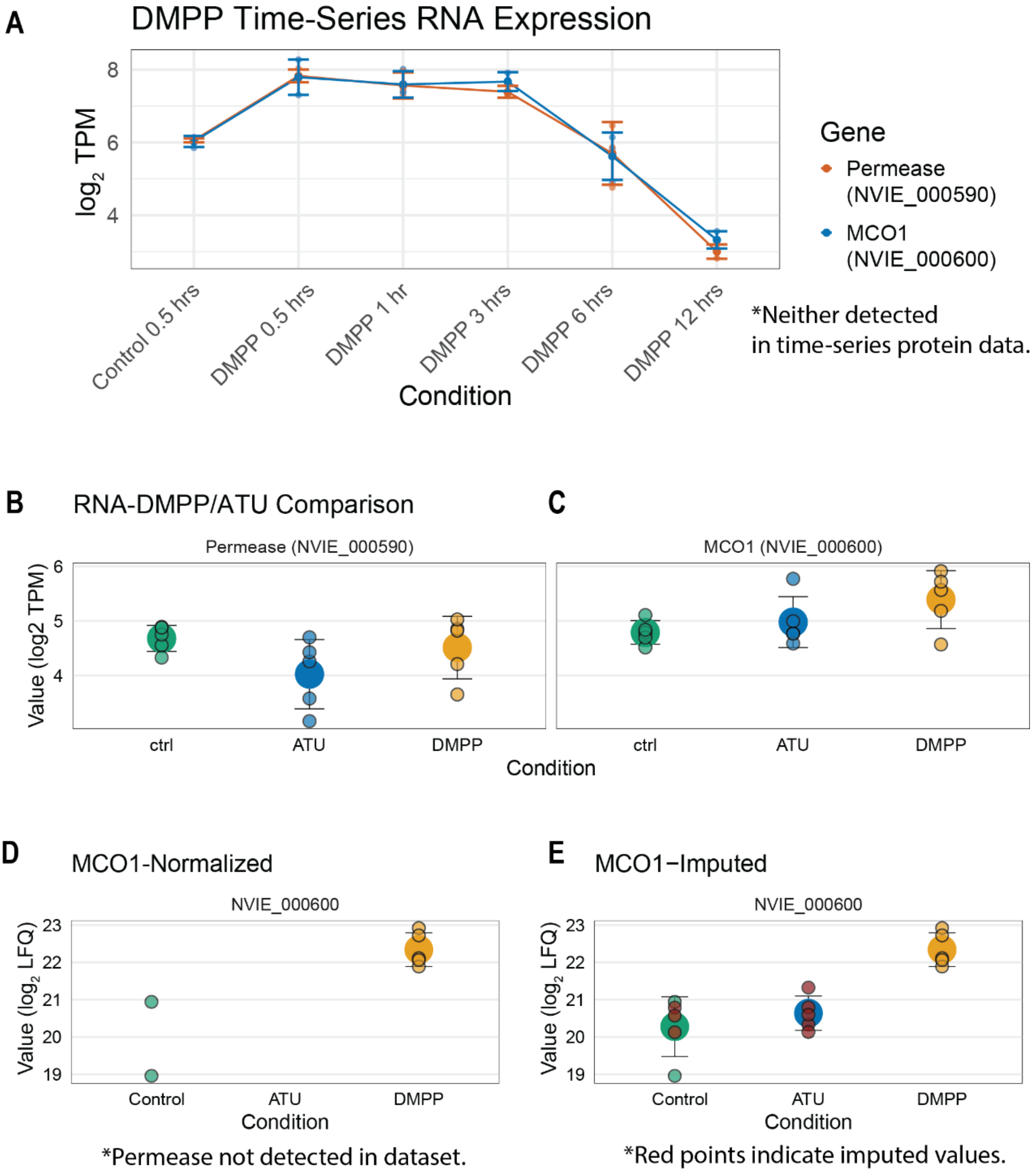


**Figure S2. Transcriptomic and proteomic abundances of multicopper oxidase MCO1 and associated permease. A)** Abundance of transcripts following addition of 2 mM DMPP. **B, C)** Transcriptomic abundance of MCO1 **(B)** and permease **(C)** in comparative DMPP and ATU experiment. **D)** Proteomic abundance of normalized MCO1 in comparative DMPP and ATU experiment. **E)** Proteomic abundance of MCO1 in comparative DMPP and ATU experiment after imputation of missing proteins. Imputed data points are marked in red. In **(B, C, D, E)** the large central point represents the average of detected or detected/imputed data points. Smaller points represent individual samples. In all plots, error bars represent standard deviation of the mean.

**A** Top 10 Up-Regulated Transcripts

ATU

| Gene Name | ATU $log_2FC$ | Protein Product |
|---|---|---|
| NME_016460 | 8.06 | conserved membrane protein of unknown function |
| NME_014210 | 6.48 | protein of unknown function |
| NME_001880 | 6.43 | protein of unknown function |
| NME_014220 | 6.31 | membrane protein of unknown function |
| NME_028550 | 5.90 | protein of unknown function |
| NME_013170 | 5.89 | protein of unknown function |
| copA2 | 5.35 | copper-exporting P-type ATPase A |
| tfb4 | 5.12 | transcription initiation factor IIB |
| NME_027670 | 5.07 | protein of unknown function |
| NME_017180 | 5.06 | hypothetical protein |

DMPP

| Gene Name | DMPP $log_2FC$ | Protein Product |
|---|---|---|
| NME_016460 | 8.40 | conserved membrane protein of unknown function |
| copA2 | 6.93 | copper-exporting P-type ATPase A |
| NME_028550 | 6.33 | protein of unknown function |
| NME_001880 | 5.89 | protein of unknown function |
| NME_027670 | 5.45 | protein of unknown function |
| NME_013170 | 5.43 | protein of unknown function |
| NME_012910 | 5.34 | putative heavy metal transport/detoxification protein |
| copA1 | 5.20 | Copper-exporting P-type ATPase A |
| tfb4 | 5.14 | transcription initiation factor IIB |
| NME_014210 | 5.09 | protein of unknown function |

**B** Top 10 Down-Regulated Transcripts

ATU

| Gene Name | ATU $log_2FC$ | Protein Product |
|---|---|---|
| NME_019240 | -7.72 | protein of unknown function |
| NME_019250 | -7.04 | Multicopper oxidase type 3 |
| NME_1975 | -6.66 | protein of unknown function |
| aniA | -5.81 | multicopper oxidase type 3 |
| NME_017740 | -5.19 | Antibiotic biosynthesis monooxygenase |
| NME_017960 | -4.35 | Carbonate dehydratase |
| hsp20-2 | -3.60 | heat-shock protein Hsp20 |
| NME_017760 | -3.29 | protein of unknown function |
| NME_015670 | -3.25 | exported protein of unknown function |
| hyfB | -3.16 | putative Hydrogenase-4 membrane subunit HyfB |

DMPP

| Gene Name | DMPP $log_2FC$ | Protein Product |
|---|---|---|
| NME_009230 | -6.22 | hypothetical protein |
| NME_021840 | -2.26 | putative Transcriptional regulator |
| NME_026950 | -2.24 | multicopper oxidase type 3 |
| lysX2 | -2.20 | lysine biosynthesis enzyme |
| NME_003800 | -2.17 | hypothetical protein |
| NME_007740 | -2.14 | putative transcriptional regulator, AsnC family |
| NME_021830 | -2.12 | protein of unknown function |
| arsB | -2.02 | putative arsenical pump membrane protein |
| NME_003690 | -2.01 | TRAM domain-containing protein |
| NME_021810 | -1.97 | protein of unknown function |

**C** Up- and Down-Regulated Proteins

ATU

| Gene Name | ATU $log_2FC$ | Protein.Product |
|---|---|---|
| NME_029950 | 3.34 | conserved protein of unknown function |
| glnA | 2.48 | glutamate-ammonia ligase / Glutamine synthetase |
| NME_017000 | 2.16 | conserved protein of unknown function |
| eIF-2B | 1.38 | translation initiation factor 2, subunit beta |
| NME_023790 | 0.94 | hypothetical protein |
| NME_008480 | 0.75 | putative dihydrolipoamide dehydrogenase |
| NME_012780 | 0.70 | Swiveling domain associated with predicted aconitase |
| NME_000010 | 0.46 | hypothetical protein |
| NME_029640 | -1.00 | voltage-gated potassium channel beta subunit family protein |
| NME_015070 | -1.02 | putative exonuclease of the beta-lactamase fold |
| NME_013080 | -1.15 | hypothetical protein |
| NME_027600 | -1.18 | putative polyketide cyclase/dehydrase |
| NME_025490 | -1.27 | transcriptional regulator, TrmB |
| NME_007040 | -1.46 | Methyltransferase type 12 |
| cheB | -1.63 | Chemotaxis response regulator protein-glutamate methylesterase |
| lysA | -1.88 | diaminopimelate decarboxylase LysA |
| NME_007540 | -2.05 | radical SAM domain protein |
| polC | -2.74 | DNA polymerase II large subunit |

DMPP

| Gene Name | DMPP $log_2FC$ | Protein.Product |
|---|---|---|
| ureC1 | 3.29 | urease subunit alpha |
| NME_029950 | 2.98 | conserved protein of unknown function |
| glnA | 2.39 | glutamate-ammonia ligase / Glutamine synthetase |
| NME_017000 | 2.17 | conserved protein of unknown function |
| hsp20-3 | 1.79 | heat-shock protein Hsp22 |
| NME_012560 | 1.57 | putative PAPS reductase |
| NME_029750 | 1.37 | hypothetical protein |
| degP | 0.96 | Protease Do-like 1, chloroplastic |
| NME_008480 | 0.92 | putative dihydrolipoamide dehydrogenase |
| NME_023790 | 0.89 | hypothetical protein |
| NME_022130 | 0.73 | exported protein of unknown function |
| NME_000010 | 0.58 | hypothetical protein |
| NME_027600 | -0.80 | putative polyketide cyclase/dehydrase |
| NME_006870 | -1.19 | putative AAA+ ATPase associated with various cellular activities |
| cheB | -1.77 | Chemotaxis response regulator protein-glutamate methylesterase |
| NME_007540 | -1.86 | radical SAM domain protein |
| lysA | -1.87 | diaminopimelate decarboxylase LysA |
| NME_000340 | -2.03 | thiosulfate sulfurtransferase |
| NME_009790 | -3.22 | putative Methyl-accepting chemotaxis sensory transducer |

**Figure S3. Up- and down-regulated transcripts and proteins in comparative DMPP and ATU experiment. A)** Top 10 up-regulated genes in ATU and DMPP treated cultures with respect to control. **B)** Top 10 down-regulated genes in ATU and DMPP treated cultures with respect to control. **C)** All up- and down-regulated genes in ATU and DMPP treated cultures with respect to control. Proteins shaded orange indicate proteins where at least one condition relied on highly on imputation (3 or more missing values). All transcriptomic data can be found in Dataset S2. All proteomic data can be found in Dataset S3.

| Cluster | Locus Tag | Gene Name | RefSeq Locus Tag | DMPP vs. control $log_2FC$ | ATU vs. control $log_2FC$ | Protein Product |
|---|---|---|---|---|---|---|
| Complex I Gene Cluster | NVE_011580 | nuoN | NVE_RS05590 | -1.84 | -2.89 | NADH-quinone oxidoreductase subunit N |
| | NVE_011590 | nuoL | NVE_RS05595 | -1.83 | -3.01 | NADH-quinone oxidoreductase subunit L |
| | NVE_011600 | nuoM | NVE_RS05600 | -1.72 | -2.65 | NADH-quinone oxidoreductase subunit M |
| | NVE_011610 | nuoK | NVE_RS05605 | -1.42 | -2.52 | NADH-quinone oxidoreductase subunit K |
| | NVE_011620 | nuoJ | NVE_RS05610 | -1.60 | -2.87 | NADH-quinone oxidoreductase subunit J |
| | NVE_011630 | nuoI | NVE_RS05615 | -1.17 | -2.09 | NADH-quinone oxidoreductase subunit I |
| | NVE_011640 | nuoH | NVE_RS05620 | -1.33 | -1.95 | NADH-quinone oxidoreductase subunit H |
| | NVE_011650 | nuoD | NVE_RS05625 | -0.58 | -0.53 | NADH-quinone oxidoreductase subunit D |
| | NVE_1185 | nuoC | NVE_RS05630 | 0.79 | 0.92 | NADH-quinone oxidoreductase subunit C |
| | NVE_011660 | nuoB | NVE_RS05635 | 1.35 | 1.60 | NADH-quinone oxidoreductase subunit B |
| | NVE_011670 | nuoA | NVE_RS05640 | -0.88 | -1.46 | NADH-quinone oxidoreductase subunit A |
| Complex V Gene Cluster | NVE_022890 | atpE | NVE_RS10985 | -1.01 | -1.27 | archaeal A1A0-type ATP synthase, subunit E |
| | NVE_022900 | atpA | NVE_RS10990 | -1.06 | -1.34 | archaeal A1A0-type ATP synthase, subunit A |
| | NVE_022910 | atpB | NVE_RS10995 | -1.07 | -1.49 | archaeal A1A0-type ATP synthase, subunit B |
| | NVE_022920 | atpD | NVE_RS11000 | -1.02 | -1.69 | archaeal A1A0-type ATP synthase, subunit D |
| | NVE_022930 | NVE_022930 | NVE_RS16155 | 0.57 | 1.07 | protein of unknown function |
| | NVE_022940 | atpK | NVE_RS11005 | -1.63 | -1.55 | archaeal A1A0-type ATP synthase, subunit K |
| | NVE_022970 | atpI | NVE_RS11010 | -1.08 | -1.27 | archaeal A1A0-type ATP synthase, subunit I |
| | NVE_022980 | NVE_022980 | NVE_RS11015 | -0.89 | -0.68 | conserved protein of unknown function |
| | NVE_022990 | atpC | NVE_RS11020 | -0.27 | -0.03 | archaeal A1A0-type ATP synthase, subunit C |

**Figure S4. Differential expression of Complex I and Complex V in comparative DMPP and ATU experiment.** Additional data can be found in Dataset S2.

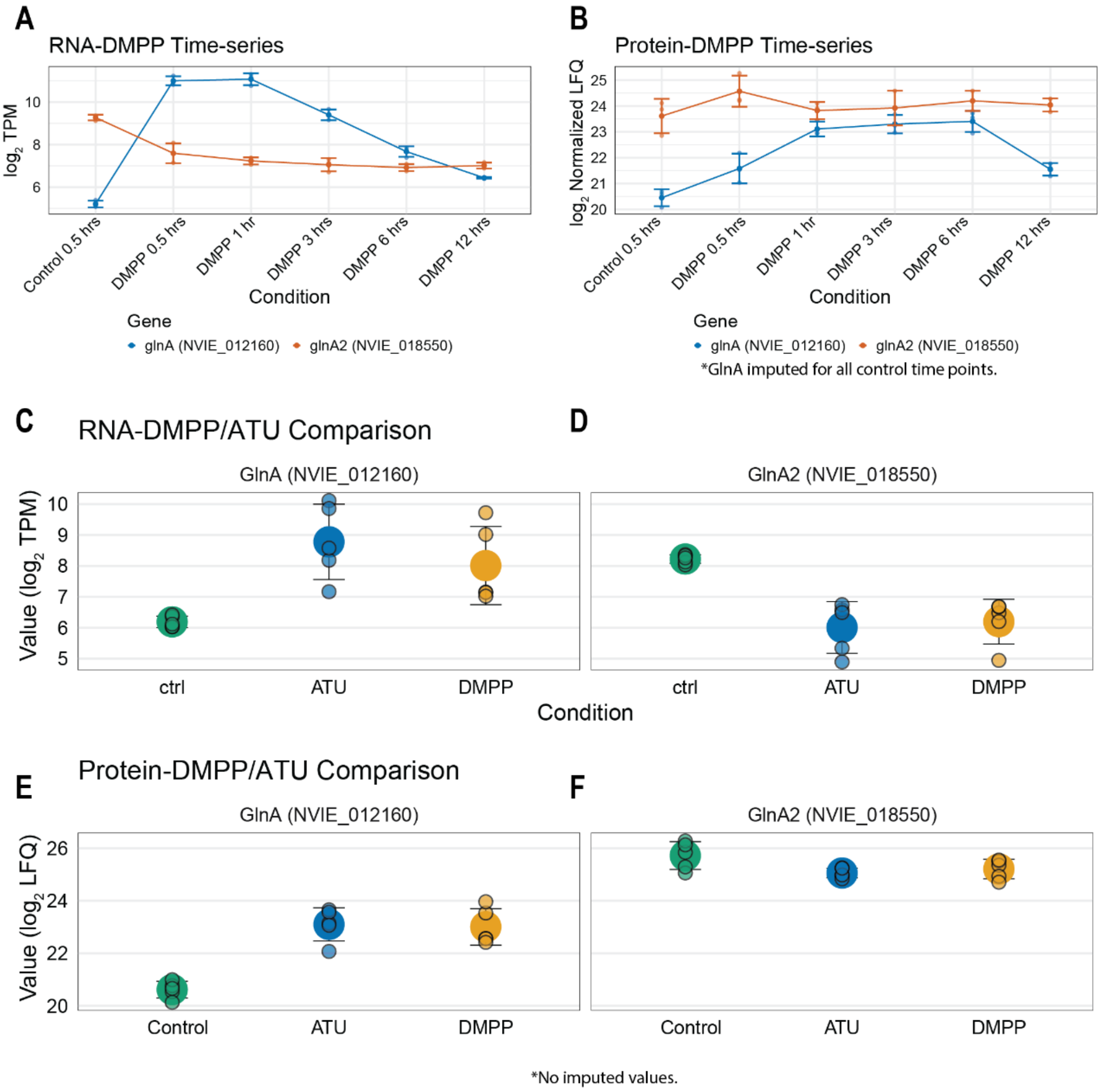


**Figure S5. Transcriptomic and proteomic abundances of glutamine synthetases (glnA and glnA2). A)** Abundance of transcripts following addition of 2 mM DMPP. **B)** Abundance of proteins following addition of 2 mM DMPP. **C,D)** Transcriptomic abundance of *glnA* **(C)** and *glnA2* **(D)** in comparative DMPP and ATU experiment. **E)** Proteomic abundance of normalized GlnA in comparative DMPP and ATU experiment. **F)** Proteomic abundance of normalized GlnA2 in comparative DMPP and ATU experiment. In **(C, D, E, F)** the large central point represents the average of detected or detected/imputed data points. Smaller points represent individual samples. In all plots, error bars represent standard deviation of the mean.

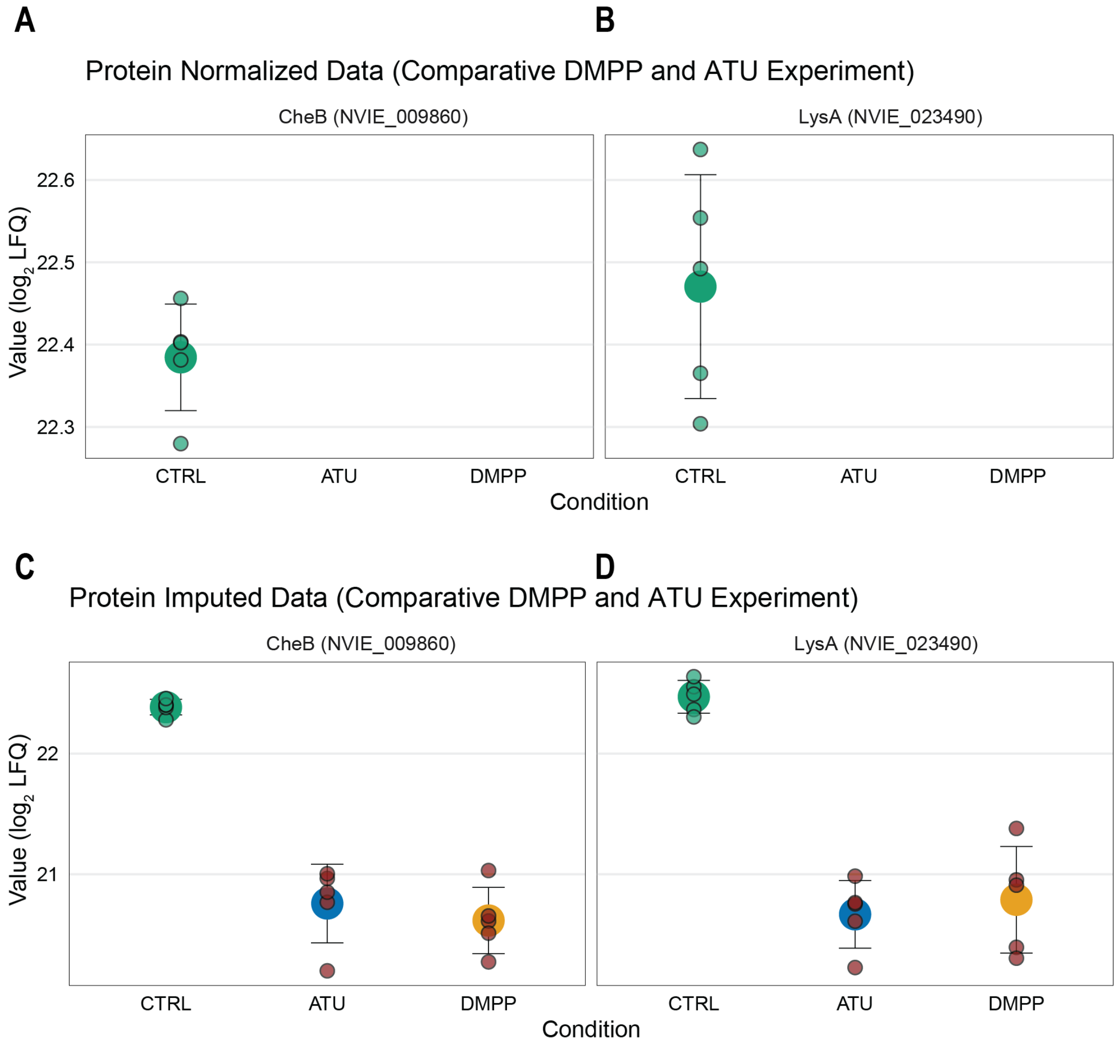


*Red points indicate imputed values.

**Figure S6. Proteomic abundances of CheB and LysA in comparative DMPP and ATU experiment. A)** Proteomic abundance of normalized CheB in comparative DMPP and ATU experiment. **B)** Proteomic abundance of normalized LysA in comparative DMPP and ATU experiment. **C)** Proteomic abundance of CheB in comparative DMPP and ATU experiment after imputation of missing proteins. **D)** Proteomic abundance of LysA in comparative DMPP and ATU experiment after imputation of missing proteins. Imputed data points are marked in red. In all panels, the large central point represents the average of detected or detected/imputed data points. Smaller points represent individual samples. In all plots, error bars represent standard deviation of the mean.

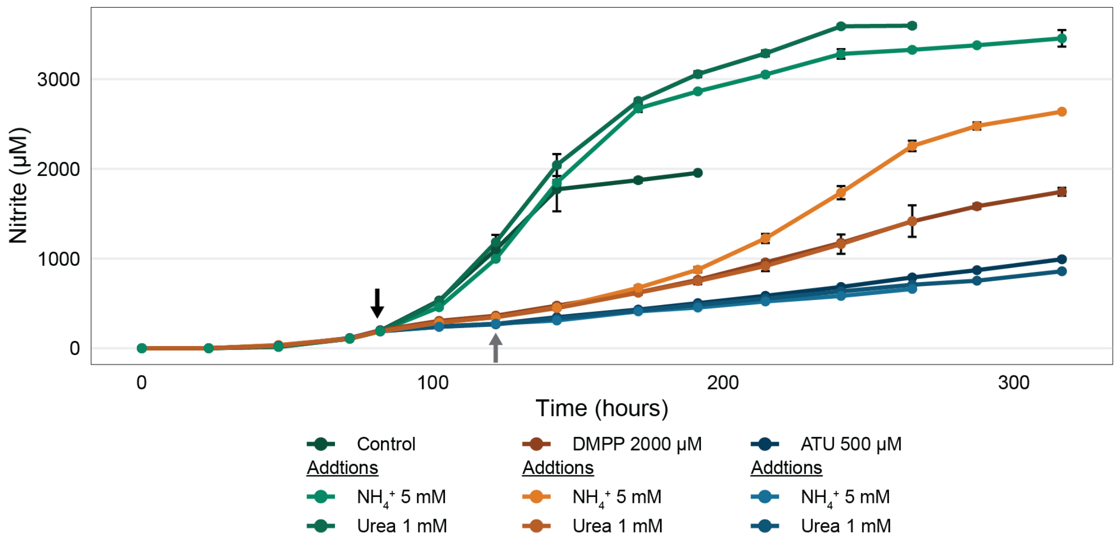


**Figure S7. Stress relief experiments *on N. viennensis* with 1 mM urea.** Additions of ammonia (5 mM) and urea (1 mM) to cultures treated with 2 mM DMPP or 0.5 mM ATU. Error bars represent standard deviations of the mean (triplicates). Black arrows indicate the time of addition of DMPP or ATU to treated cultures and addition of ammonium or urea to control cultures. Grey arrows represent time of additions of ammonium or urea to cultures treated with DMPP or ATU. Cultures with DMPP and ammonium served as a positive control. No relief was observed in DMPP with 1 mM of urea or in ATU with either 5 mM ammonium or 1 mM urea.

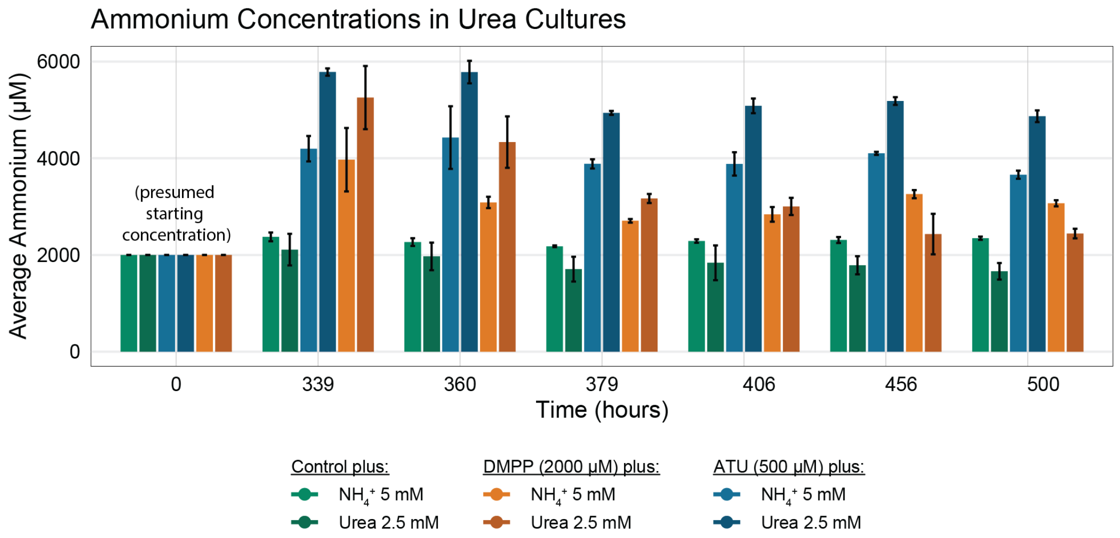


**Figure S8. Ammonium measurements in cultures with additional ammonium and urea.** Colored bars represent the average amount of detected ammonium. Error bars represent the standard deviation of the mean. Ammonium levels correspond to growth curves in Figure 7.

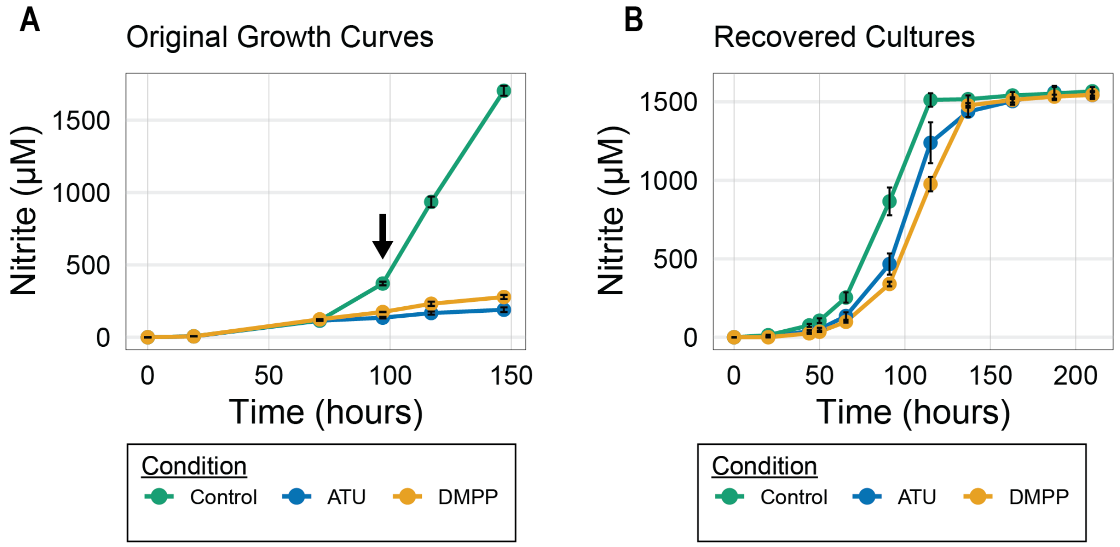


**Figure S9. Recovery of cultures of *N. viennensis* from exposure to 2 mM DMPP or 0.5 mM ATU. A)** *N. viennensis* cultures grown with 2 mM DMPP or 0.5 mM ATU or without inhibitors. Black arrow indicates the point where samples were taken for subsequent washing and inoculation to fresh medium (see supplementary materials and methods). **B)** Washed and reinoculated cultures derived from the treated cultures shown in panel A. Error bars represent the standard deviation of the mean.

| | $log_2FC$ | | | | |
|---|---|---|---|---|---|
| | **Cu lim** | **DMPP** | **ATU** | **Gene Name** | **Protein Product** |
| Up-regulated under copper limiation | 6.99 | -1.22 | -7.04 | NVE_019250 | Multicopper oxidase type 3 |
| | 5.53 | 4.15 | 4.11 | NVE_001850 | exported protein of unknown function |
| | 4.75 | 5.09 | 6.48 | NVE_014210 | protein of unknown function |
| | 4.59 | 4.98 | 6.31 | NVE_014220 | membrane protein of unknown function |
| | 4.39 | 1.49 | 2.04 | NVE_018920 | protein of unknown function |
| | 4.27 | -1.21 | -5.81 | aniA | multicopper oxidase type 3 |
| | 4.24 | 1.64 | 1.76 | NVE_1990 | protein of unknown function |
| | 4.23 | 2.18 | 2.53 | NVE_1444 | hypothetical protein |
| | 4.15 | -0.73 | -0.42 | NVE_012190 | protein of unknown function |
| | 4.11 | 2.92 | 3.07 | NVE_024190 | protein of unknown function with C-terminal blue (Type 1) copper domain |
| | 3.98 | 1.52 | 2.48 | NVE_014170 | putative sialidase - neuraminidase family protein |
| | 3.75 | 8.40 | 8.06 | NVE_016460 | conserved membrane protein of unknown function |
| | 3.54 | 0.52 | 0.56 | NVE_026610 | conserved exported protein of unknown function |
| | 3.50 | NA | NA | NVE_018430 | protein of unknown function |
| | 3.49 | 1.08 | 0.64 | NVE_023360 | exported protein of unknown function |
| | 3.33 | -1.53 | -7.72 | NVE_019240 | protein of unknown function |
| | 3.24 | 1.13 | 2.45 | NVE_012970 | protein of unknown function |
| | 3.20 | -0.39 | 0.06 | NVE_000930 | putative peptidase M10A |
| | 3.20 | 0.19 | -3.29 | NVE_017760 | protein of unknown function |
| | 3.13 | 0.40 | 0.55 | NVE_026600 | protein of unknown function |
| | 3.13 | 2.23 | 0.83 | NVE_001840 | putative Zn-finger domain containing protein |
| | 3.03 | 0.74 | 1.85 | NVE_012980 | Radical SAM |
| | 3.03 | 0.50 | 0.52 | NVE_016680 | putative exported polysaccharide deacetylase family protein |
| | 2.99 | 0.23 | 1.52 | NVE_013000 | protein of unknown function |
| | 2.97 | -0.42 | -6.66 | NVE_1975 | protein of unknown function |
| Down-regulated under copper limitation | -3.51 | -0.17 | -0.53 | NVE_001000 | putative surface-associated Ca2+-binding protein, Haemolysin-type |
| | -2.32 | -0.59 | -1.27 | NVE_017190 | putative transposase IS605 OrfB family |
| | -2.01 | -1.73 | -2.40 | NVE_021040 | conserved protein of unknown function |
| | -1.81 | -2.01 | -2.48 | NVE_003690 | TRAM domain-containing protein |
| | -1.73 | -1.37 | -1.38 | NVE_028970 | protein of unknown function |
| | -1.67 | 1.33 | 1.38 | NVE_010050 | protein of unknown function |
| | -1.65 | -0.25 | -0.02 | NVE_000990 | hypothetical protein |
| | -1.64 | -0.85 | -2.11 | NVE_002250 | putative cupredoxin |
| | -1.63 | 0.10 | 1.13 | NVE_019730 | protein of unknown function |
| | -1.60 | 0.81 | -3.98 | NVE_006760 | protein of unknown function |
| | -1.60 | 0.24 | 0.61 | NVE_001020 | putative cell wall surface anchor family protein |
| | -1.59 | -0.89 | -2.11 | NVE_002270 | protein of unknown function |
| | -1.55 | -1.10 | -2.40 | NVE_025500 | SNF7-domain-containing protein |
| | -1.55 | -1.03 | -1.20 | NVE_022410 | Rieske (2Fe-2S) domain-containing protein |
| | -1.52 | -1.58 | -1.79 | NVE_005290 | protein of unknown function |
| | -1.47 | -0.51 | -1.14 | NVE_016390 | exported protein of unknown function |
| | -1.47 | -0.71 | -1.42 | NVE_002700 | hypothetical protein |
| | -1.42 | 0.29 | -0.48 | cimA | putative 2-isopropylmalate synthase |
| | -1.41 | 1.90 | 0.33 | NVE_021540 | protein of unknown function |
| | -1.40 | -1.24 | -0.69 | NVE_007060 | putative NADH-quinone oxidoreductase, subunit D - related |
| | -1.39 | 3.95 | -0.32 | NVE_017620 | protein of unknown function |
| | -1.38 | 0.06 | -0.09 | NVE_001320 | hypothetical protein |
| | -1.38 | 1.76 | 1.82 | NVE_000940 | cell surface protein with DUF11 domain |
| | -1.38 | 1.13 | 1.30 | NVE_019740 | protein of unknown function |
| | -1.37 | -1.32 | -1.92 | NVE_028050 | ABC uptake transporter, substrate-binding protein |

**Figure S10. Comparison of genes in *N. viennensis* that are highly responsive to copper limitation.** The top 25 up- and down-regulated genes from the study by Reyes et al (2020)[33] are shown along with the same genes and their response to 2 mM DMPP or 0.5 mM ATU. Values shaded grey indicate genes that did not pass the adjusted *P* value cut-off of 0.001.

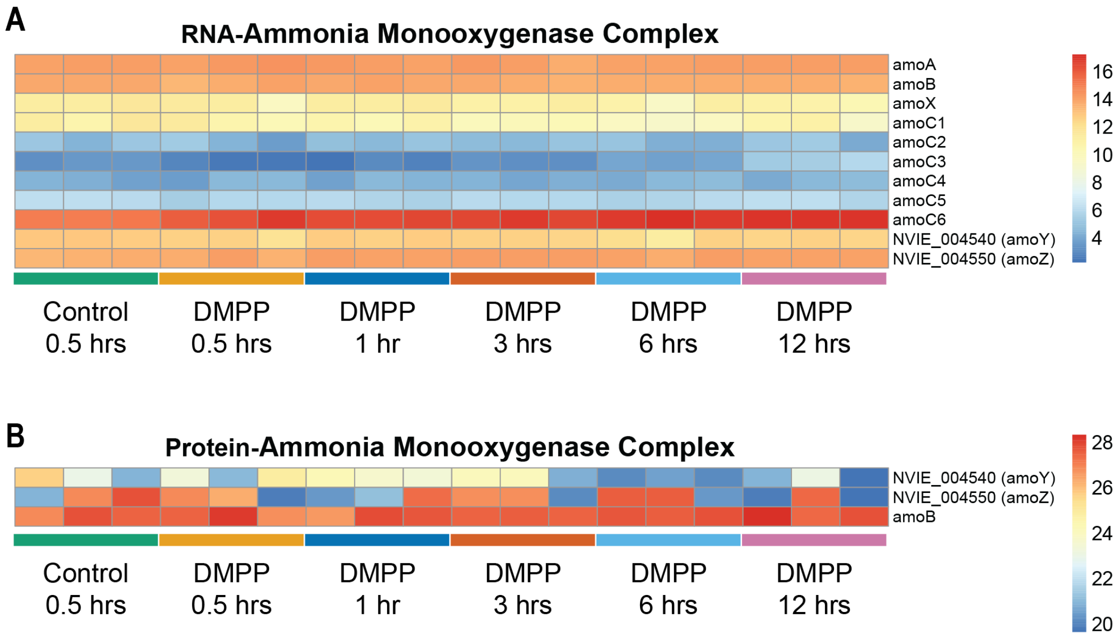


**Figure S11. Response of the ammonia monooxygenase genes (A) and proteins (B) during the course of the DMPP time-series experiment.**

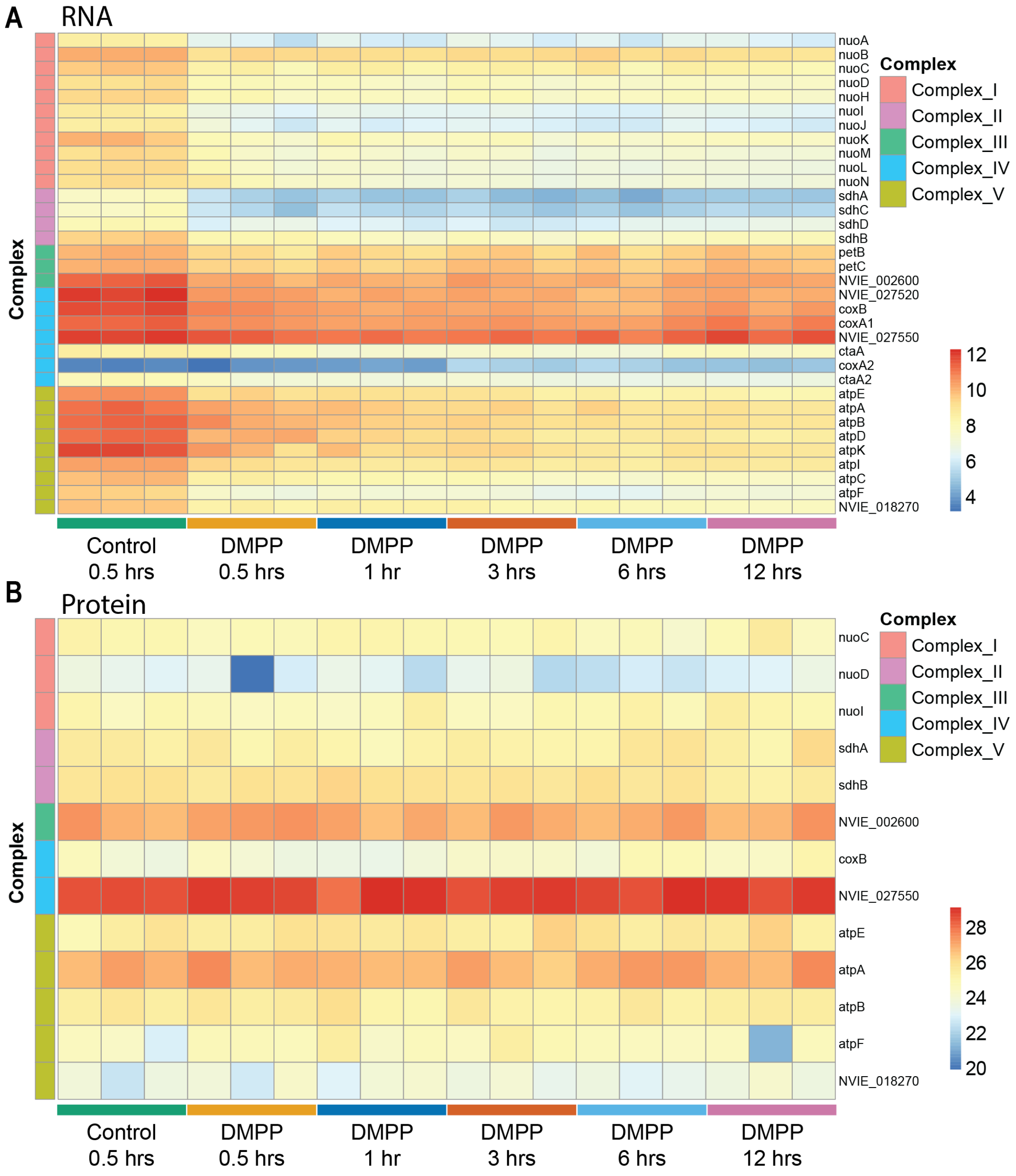


**Figure S12. Response of electron transport chain complex genes (A) and proteins (B) during the course of the DMPP time-series experiment.**

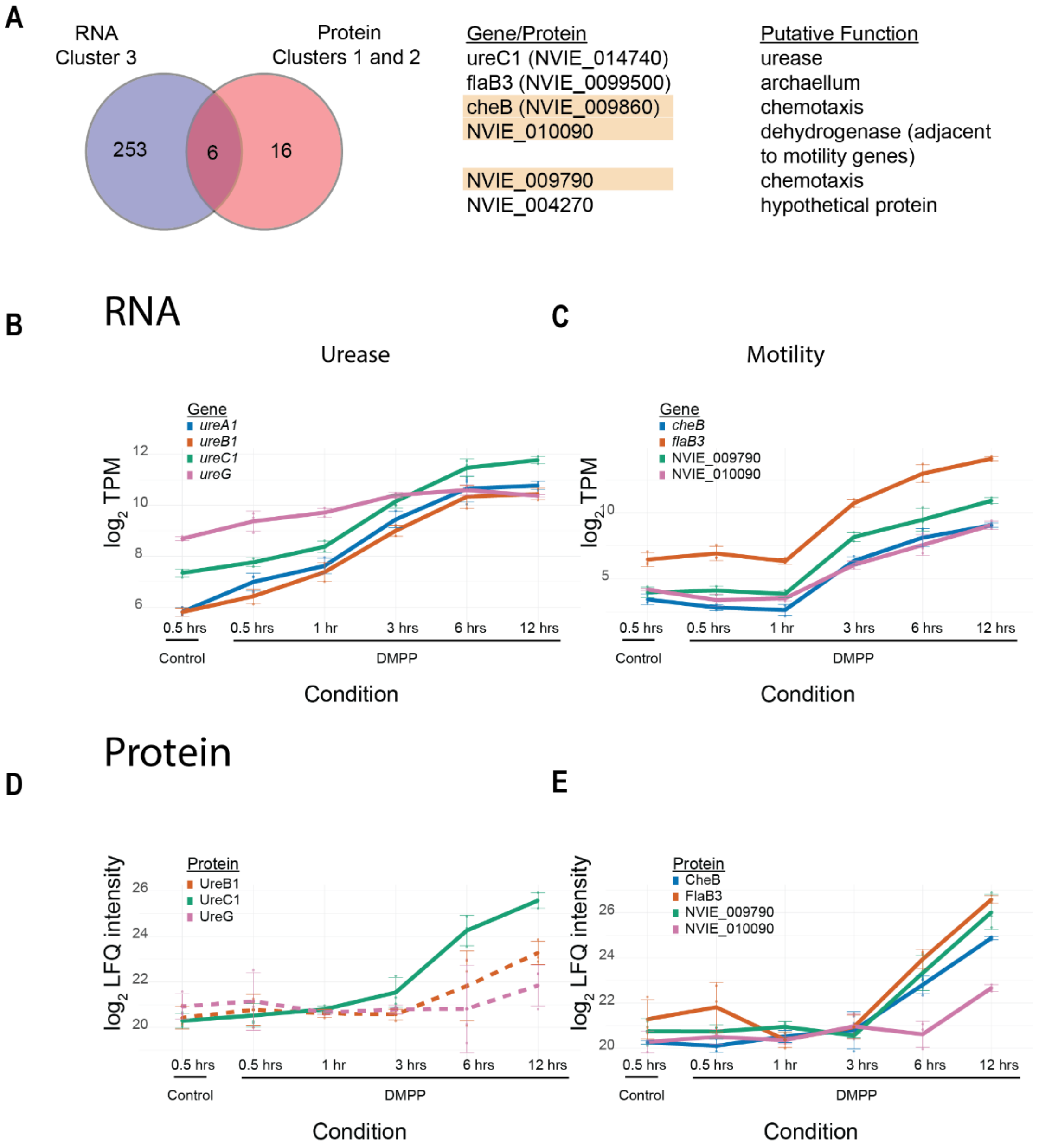


**Figure S13. A)** Venn diagram between the RNA cluster 3 (RNA with increasing relative expression) and protein clusters 1 and 2 (proteins with increasing relative abundance), with 6 genes showing the same trend at both transcriptomic and proteomic level. Genes/proteins with an orange background indicate proteins where 67% or more of replicates relied on imputation due to missing values. **(B, C)** RNA expression is presented as $log_2$-transformed Transcripts Per Million (log2 TPM), **(D, E)** protein levels are presented as $log_2$-transformed LFQ intensities. Individual data points represent biological triplicates, solid lines indicate the mean, and error bars represent the standard deviation. Colors correspond to specific gene/protein targets as defined in the legend. **B)** RNA expression of main urease operon and ureG, **C)** RNA expression of selected motility genes, **D)** protein levels of main urease operon and ureG, with dotted line are denoted proteins that were not statistically significant based on ANOVA ($p < 0.05$) but present in the proteome and **E)** protein levels of selected motility genes.

### Motility Gene Cluster Expression Comparison

Average of log2 TPM for three replicates in each condition of Time series and for five replicates for the DMPP vs ATU exp.

| Locus Tag | Gene Name | Time-series Control | DMPP/ATU Comparative Experiment Control | DMPP 0.5 hrs | DMPP 1 hr | DMPP 3 hrs | DMPP 6 hrs | DMPP 12 hrs | DMPP | ATU | Protein Product |
|---|---|---|---|---|---|---|---|---|---|---|---|
| NME_009600 | flaK1 | 4.81 | 1.75 | 5.14 | 5.38 | 5.39 | 5.49 | 5.51 | 1.37 | 1.78 | putative archaeal preflagellin peptidase FlaK |
| NME_009610 | NME_009610 | 3.77 | 2.05 | 3.40 | 3.85 | 4.01 | 4.01 | 4.05 | 2.37 | 0.59 | hypothetical protein |
| NME_009620 | NME_009620 | 0.00 | 0.00 | 0.00 | 0.00 | 0.00 | 0.00 | 0.00 | 0.00 | 0.00 | protein of unknown function |
| NME_009630 | NME_009630 | 6.21 | 3.03 | 5.46 | 5.42 | 5.57 | 5.41 | 5.86 | 1.43 | 2.02 | zinc finger C2H2 domain-containing protein |
| NME_009640 | NME_009640 | 2.12 | 0.62 | 2.07 | 2.43 | 2.13 | 2.14 | 2.09 | 1.36 | 1.35 | protein of unknown function |
| NME_009650 | NME_009650 | 8.30 | 2.73 | 7.64 | 7.92 | 7.87 | 7.82 | 8.01 | 2.22 | 2.64 | putative integrase family protein |
| NME_009660 | NME_009660 | 1.47 | 0.00 | 1.46 | 1.43 | 1.35 | 1.43 | 1.49 | 0.00 | 0.65 | protein of unknown function |
| NME_0991 | NME_0991 | 0.00 | 0.15 | 0.00 | 0.00 | 0.00 | 0.00 | 0.00 | 0.00 | 0.00 | protein of unknown function |
| NME_009670 | NME_009670 | 2.86 | 2.14 | 3.07 | 3.17 | 3.21 | 3.05 | 3.62 | 4.35 | 2.68 | protein of unknown function |
| NME_009680 | NME_009680 | -1.59 | 0.00 | -1.63 | -1.63 | -1.63 | -1.63 | -1.63 | 0.00 | 0.00 | protein of unknown function |
| NME_009690 | NME_009690 | 2.81 | 0.73 | 3.04 | 3.30 | 2.89 | 2.68 | 2.89 | 0.00 | 0.78 | protein of unknown function |
| NME_009700 | NME_009700 | 1.41 | 0.00 | 1.35 | 1.32 | 1.41 | 1.86 | 1.44 | 0.00 | 0.81 | protein of unknown function |
| NME_009710 | NME_009710 | 0.00 | 0.69 | 0.00 | 0.00 | 0.00 | 0.00 | 0.00 | 0.00 | 0.00 | protein of unknown function |
| NME_009720 | NME_009720 | 8.19 | 4.54 | 7.07 | 6.69 | 6.94 | 7.05 | 7.31 | 3.65 | 3.72 | protein of unknown function |
| NME_009730 | NME_009730 | 11.62 | 7.62 | 11.88 | 11.83 | 11.86 | 11.80 | 11.81 | 7.77 | 7.84 | uncharacterised protein |
| NME_009740 | NME_009740 | 10.29 | 7.91 | 9.84 | 9.75 | 9.76 | 9.59 | 9.95 | 7.00 | 6.65 | putative nucleic acid binding OB-fold tRNA/helicase-type |
| NME_009750 | NME_009750 | 6.45 | 5.31 | 6.87 | 7.10 | 7.10 | 7.05 | 6.96 | 7.05 | 7.04 | protein of unknown function |
| NME_009760 | NME_009760 | 10.50 | 7.40 | 10.14 | 9.99 | 10.02 | 9.96 | 10.27 | 7.22 | 7.01 | putative protein of unknown function DUF192 |
| NME_009770 | flaB1 | 8.27 | 5.89 | 8.28 | 8.26 | 9.05 | 10.16 | 12.21 | 4.33 | 4.50 | archaeal flagellin |
| NME_009780 | NME_009780 | 9.02 | 4.58 | 8.65 | 8.54 | 10.25 | 11.51 | 12.87 | 3.37 | 3.17 | putative transcriptional regulator, TrmB |
| NME_009790 | NME_009790 | 11.39 | 7.46 | 11.62 | 11.50 | 13.71 | 14.76 | 15.95 | 6.09 | 5.90 | putative Methyl-accepting chemotaxis sensory transducer |
| NME_009800 | NME_009800 | 7.18 | 5.91 | 6.80 | 6.83 | 6.89 | 6.77 | 6.81 | 5.06 | 4.11 | protein of unknown function |
| NME_009810 | NME_009810 | 8.85 | 6.98 | 8.90 | 9.10 | 9.16 | 9.04 | 9.04 | 6.27 | 6.55 | protein of unknown function |
| NME_009820 | NME_009820 | 8.54 | 4.99 | 8.19 | 8.17 | 9.80 | 11.05 | 12.51 | 4.05 | 3.90 | putative methyl-accepting chemotaxis protein |
| NME_009840 | cheW | 8.62 | 7.39 | 8.39 | 8.29 | 10.40 | 11.60 | 12.46 | 5.35 | 5.10 | chemotaxis protein CheW |
| NME_009850 | cheY | 7.14 | 6.55 | 7.19 | 6.95 | 9.12 | 10.35 | 11.28 | 4.73 | 3.59 | chemotaxis response regulator CheY |
| NME_009860 | cheB | 9.03 | 6.40 | 8.99 | 8.90 | 10.84 | 12.20 | 13.01 | 4.52 | 3.86 | Chemotaxis response regulator protein-glutamate methylesterase |
| NME_009870 | cheA | 10.20 | 5.89 | 9.63 | 9.71 | 11.25 | 12.56 | 13.65 | 4.50 | 3.89 | chemotactic sensor histidine kinase CheA |
| NME_009880 | cheC | 8.92 | 7.28 | 8.88 | 8.51 | 10.09 | 11.47 | 12.60 | 5.53 | 5.10 | putative chemotaxis protein cheC |
| NME_009890 | cheD | 7.34 | 5.25 | 6.96 | 6.96 | 8.24 | 9.31 | 10.53 | 3.82 | 2.99 | chemoreceptor glutamine deamidase CheD |
| NME_009900 | cheR | 6.49 | 3.19 | 6.39 | 5.96 | 7.03 | 8.19 | 9.45 | 1.57 | 1.03 | putative chemotaxis MCP methyltransferase CheR |
| NME_009910 | flaI | 8.21 | 4.79 | 8.15 | 8.25 | 10.00 | 11.18 | 12.23 | 3.29 | 2.80 | archaeal flagella protein FlaI |
| NME_009920 | trmB1 | 10.99 | 8.00 | 11.31 | 11.67 | 13.07 | 13.59 | 14.76 | 6.52 | 6.09 | putative transcriptional regulator, TrmB |
| NME_009930 | flaB2 | 10.57 | 6.77 | 11.18 | 11.27 | 10.96 | 11.05 | 10.78 | 5.32 | 4.08 | Archaeal flagellin |
| NME_009950 | flaB3 | 12.32 | 10.55 | 12.61 | 12.41 | 14.44 | 16.13 | 17.10 | 8.90 | 8.60 | archaeal flagellin |
| NME_009960 | NME_009960 | 7.56 | 3.53 | 7.77 | 7.52 | 9.46 | 10.92 | 12.02 | 2.15 | 2.17 | protein of unknown function |
| NME_009970 | flaG | 7.26 | 5.15 | 6.89 | 6.74 | 8.50 | 9.81 | 11.09 | 3.45 | 3.20 | putative Archaeal flagellar protein FlaG |
| NME_009980 | flaF | 7.45 | 5.22 | 7.22 | 7.17 | 8.67 | 9.98 | 11.27 | 4.09 | 3.51 | putative flagellar protein FlaF |
| NME_009990 | flaH | 7.63 | 4.88 | 7.60 | 7.31 | 8.68 | 9.83 | 11.03 | 3.24 | 3.27 | flagella protein FlaH |
| NME_010000 | flaJ | 7.94 | 3.99 | 7.89 | 7.95 | 9.10 | 10.36 | 11.52 | 2.96 | 2.48 | putative archaeal flagella assembly protein J |
| NME_010010 | NME_010010 | 6.83 | 6.58 | 7.13 | 6.99 | 9.07 | 10.36 | 11.33 | 5.42 | 4.41 | protein of unknown function |
| NME_010020 | NME_010020 | 7.54 | 7.02 | 7.94 | 7.66 | 9.66 | 11.23 | 12.10 | 6.29 | 5.51 | protein of unknown function |
| NME_010030 | NME_010030 | 10.04 | 8.73 | 10.42 | 10.22 | 12.41 | 13.81 | 14.78 | 6.99 | 6.63 | putative KaiC family protein kinase |
| NME_010040 | NME_010040 | 7.31 | 5.75 | 7.57 | 7.44 | 9.55 | 11.02 | 12.01 | 4.06 | 3.73 | protein of unknown function |
| NME_010050 | NME_010050 | 7.86 | 3.00 | 8.52 | 8.60 | 7.89 | 7.55 | 7.61 | 3.29 | 3.57 | protein of unknown function |
| NME_010060 | trmB2 | 9.65 | 5.71 | 11.22 | 11.58 | 11.54 | 11.41 | 11.16 | 5.37 | 4.48 | putative transcriptional regulator, TrmB |
| NME_010070 | NME_010070 | 4.35 | 3.03 | 4.67 | 4.68 | 4.26 | 4.38 | 4.76 | 1.61 | 1.19 | protein of unknown function |
| NME_010080 | flaK2 | 6.30 | 4.76 | 6.41 | 6.50 | 8.72 | 9.75 | 10.89 | 2.92 | 2.81 | putative archaeal preflagellin peptidase FlaK |
| NME_010090 | NME_010090 | 9.23 | 6.16 | 9.14 | 9.16 | 10.54 | 11.68 | 12.89 | 5.47 | 5.25 | Dehydrogenase |

Time-Series Experiment: DMPP 0.5 hrs, DMPP 1 hr, DMPP 3 hrs, DMPP 6 hrs, DMPP 12 hrs

DMPP/ATU Comparative Experiment: DMPP, ATU

**Figure S14. Motility gene cluster RNA expression in time-series and comparative DMPP and ATU experiment.** Full data can be found in Dataset S2.
